# Antibodies with potent and broad neutralizing activity against antigenically diverse and highly transmissible SARS-CoV-2 variants

**DOI:** 10.1101/2021.02.25.432969

**Authors:** Lingshu Wang, Tongqing Zhou, Yi Zhang, Eun Sung Yang, Chaim A. Schramm, Wei Shi, Amarendra Pegu, Olamide K. Oloninyi, Amy Ransier, Samuel Darko, Sandeep R. Narpala, Christian Hatcher, David R. Martinez, Yaroslav Tsybovsky, Emily Phung, Olubukola M. Abiona, Evan M. Cale, Lauren A. Chang, Kizzmekia S. Corbett, Anthony T. DiPiazza, Ingelise J. Gordon, Kwanyee Leung, Tracy Liu, Rosemarie D. Mason, Alexandra Nazzari, Laura Novik, Adam S. Olia, Nicole A. Doria-Rose, Tyler Stephens, Christopher D. Stringham, Chloe Adrienna Talana, I-Ting Teng, Danielle Wagner, Alicia T. Widge, Baoshan Zhang, Mario Roederer, Julie E. Ledgerwood, Tracy J. Ruckwardt, Martin R. Gaudinski, Ralph S. Baric, Barney S. Graham, Adrian B. McDermott, Daniel C. Douek, Peter D. Kwong, John R Mascola, Nancy J. Sullivan, John Misasi

**Author notes:** Equal contributions (Lingshu Wang, Tongqing Zhou, John Misasi).

## Abstract

The emergence of highly transmissible SARS-CoV-2 variants of concern (VOC) that are resistant to therapeutic antibodies highlights the need for continuing discovery of broadly reactive antibodies. We identify four receptor-binding domain targeting antibodies from three early-outbreak convalescent donors with potent neutralizing activity against 12 variants including the B.1.1.7 and B.1.351 VOCs. Two of them are ultrapotent, with sub-nanomolar neutralization titers (IC_50_ <0.0006 to 0.0102 μg/mL; IC80 < 0.0006 to 0.0251 μg/mL). We define the structural and functional determinants of binding for all four VOC-targeting antibodies, and show that combinations of two antibodies decrease the *in vitro* generation of escape mutants, suggesting potential means to mitigate resistance development. These results define the basis of therapeutic cocktails against VOCs and suggest that targeted boosting of existing immunity may increase vaccine breadth against VOCs.

**One Sentence Summary:** Ultrapotent antibodies from convalescent donors neutralize and mitigate resistance of SARS-CoV-2 variants of concern.

## Main Text

Since the start of the SARS-CoV-2 outbreak, >100 million people have been infected and >2 million have died from COVID-19 (*1*). Shortly after the first Wuhan Hu-1 (WA-1) genome sequence was published (*2*), spike proteins were generated for use in spike-specific antibody discovery (*3*–*5*). Recently, virus variants first detected in the UK (e.g., B.1.1.7)(*6*), South Africa (e.g., B.1.351) (*7*) and Brazil (P.1) (*8,9*) have been shown to contain mutations that mediate resistance to therapeutic monoclonal antibodies, have increased transmissibility and to potentially increase pathogenicity (*10*–*14*). Additionally, vaccines designed based on the original WA-1 outbreak strain sequence elicit antibody responses that show decreased *in vitro* neutralizing activity against variants (*14*–*16*). In this study, we investigated antibodies isolated from convalescent subjects who were infected by the WA-1 strain during the first few months of the outbreak, determined their reactivity against variants of concern (VOCs) and defined the structural features of their binding to spike.

We obtained blood from four mild to moderately ill WA-1-infected subjects between 30 and 50 days after symptom onset. CD19+/CD20+/IgM-/IgA+ or IgG+ B cells were sorted for binding to S-2P, receptor binding domain-subdomain-1 (RBD-SD1) or the S1 domain and individual B-cell receptors were sequenced (Figure 1A, Figure S1). In total, we sorted 889 B cells and recovered 709 (80%) paired heavy and light chain sequences and selected 200 antibodies for expression. Among the 200 antibodies, there was a broad response across all spike domains with 77 binding RBD, 46 binding N-terminal domain (NTD), 58 binding the S2 domain, and 19 binding an indeterminant epitope or failing to recognize spike in a MSD binding assay (Figure 1B). Among these, 4 RBD targeting antibodies, A19-46.1, A19-61.1, A23-58.1 and B1-182.1, were shown to have especially potent pseudovirus neutralization (IC_50_ 0.0025-0.0709 μg/mL) (Figure 1C, E). Live virus neutralization (*17*) revealed similar high potent neutralization by all four antibodies (IC_50_ 0.0021-0.0048 μg/mL) (Figure 1D-E). All antibody Fabs exhibited nanomolar affinity for SARS-CoV-2 S-2P (i.e., 2.3-7.3 nM), consistent with their potent neutralization (Figure 1E).

**Fig. 1.**
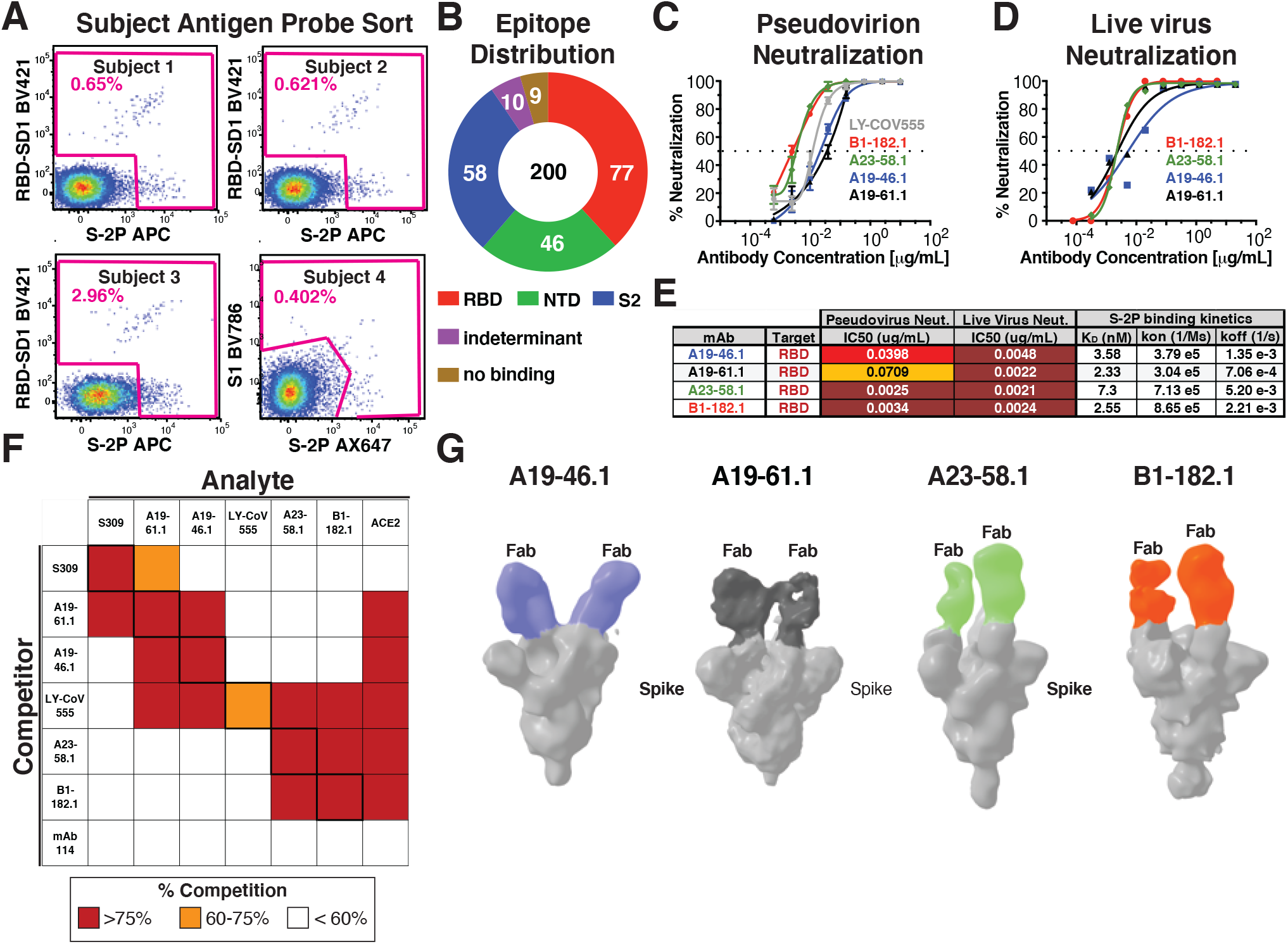
Identification and classification of highly potent antibodies from convalescent SARS-CoV-2 subjects. **(A)** Final flow cytometry sorting gate of CD19+/CD20+/IgG+ or IgA+ PBMCs for four convalescent subjects (Subjects 1-4). Shown is the staining for RBD-SD1 BV421, S1 BV786 and S-2P APC or Ax647. Cells were sorted using indicated sorting gate (pink) and percent positive cells that were either RBD-SD1, S1 or S-2P positive is shown for each subject. **(B)** Gross binding epitope distribution was determined using an MSD-based ELISA testing against RBD, NTD, S1, S-2P or HexaPro. S2 binding was inferred by S-2P or HexaPro binding without binding to other antigens. Indeterminant epitopes showed a mixed binding profile. Total number of antibodies (i.e., 200) and absolute number of antibodies within each group is shown. **(C)** Lentivirus particles pseudotyped with WA-1 spike were used to test the neutralization capacity of the indicated antibodies (n=3). **(D)** Live virus neutralization assay for A23-58.1 (n=2), A19-46.1 (n=2), A19-61.1 (n=2) and B1-182.1 (n=3). **(E)** Table showing antibody binding target, IC_50_ for pseudovirus and live virus neutralization and Fab:S-2P binding kinetics (n=2) for the indicated antibodies. **(F)** Biolayer interferometry-based epitope binning experiment. Competitor antibody (y-axis) is bound to S-2P prior to incubation with the analyte antibody or ACE2 protein (x-axis) as indicated and percent competition range bins are shown as red (>=75%), orange (60-75%) or white <60%) (n=2). mAb114 is an anti-Ebola glycoprotein antibody and is included as a negative control (*37*) **(G)** Negative stain 3D reconstructions of SARS-CoV-2 spike and Fab complexes. A19-46.1 and A19-61.1 bind to RBD in the down position while A23-58.1 and B1-182.1 bind to RBD in the up position. Representative classes were shown with 2 Fabs bound, though stoichiometry at 1 to 3 were observed.

Since VOCs have been reported to contain mutations that confer resistance to RBD-directed therapeutic antibodies such as LY-CoV555 (*18*–*20*), we examined whether the epitopes targeted by the four high-potency antibodies were distinct from LY-CoV555. We used a biolayer interferometry-based (BLI) competition binding assay to compare the binding profile of these antibodies to LY-CoV555. We noted that while LY-CoV555 prevented the binding of each of the experimental antibodies, the block was not bidirectional; the experimental antibodies did not impact the binding of LY-CoV555. This suggests that these antibodies bind distinct epitopes from LY-CoV555 (Figure 1F). We found that A23-58.1 and B1-182.1 exhibit similar binding profiles and that A19-61.1 and A19-46.1 likewise display a shared binding pattern. However, the latter two antibodies can be distinguished from each other by their capacity to compete for binding with the RBD-targeting antibody S309 (*21*) (Figure 1F). S309 binds an epitope in RBD that is accessible in the up or down position but does not compete with the SARS-CoV-2 receptor protein, angiotensin-converting enzyme (ACE2), and is a Class III RBD antibody (*22*). To further classify the antibodies, we examined whether these antibodies prevent the binding of ACE2 to spike proteins. We noted that in both BLI-competition and cell surface binding assays, all four experimental antibodies prevented the binding of ACE2 to spike (Figure 1F, Figure S2). This suggests that A19-46.1, A23-58.1 and B1-182.1 neutralize infection by blocking the interaction of RBD with ACE2 and would be classified as either Class I (i.e., ACE2 blocking, binding RBD up only) or II (i.e., ACE2 blocking, binding RBD up or down) RBD antibodies (*22*). A19-61.1 competes with S309 and blocks ACE2 binding suggesting that it may sterically block ACE2 binding similar to the Class III antibody REGN10987. To refine the classification of these antibodies, we performed negative stain 3D reconstruction and found that A19-46.1 and A19-61.1 bound near one another with RBD in the down position (Figure 1G), consistent with them being Class II and Class III antibodies, respectively. Similarly, A23-58.1 and B1-182.1 bound to overlapping regions when RBD is in the up position, suggesting that they are Class I antibodies.

Because each donor subject was infected with ancestral WA-1 variants, we evaluated antibody activity against recently emerged variants like D614G, which has become the dominant variant across the world (*23*). We observed that, similar to LY-CoV555, neutralization potency was increased against D614G compared to WA-1, with the IC_50_ and IC_80_ of each experimental antibody 1.4 to 6.3-fold lower than that seen for the WA-1 (IC_50_ of 0.0008-0.0203 μg/mL and IC_80_ of 0.0026-0.0435 μg/mL) (Figure 2A,C, Figure S3).

**Fig. 2.**
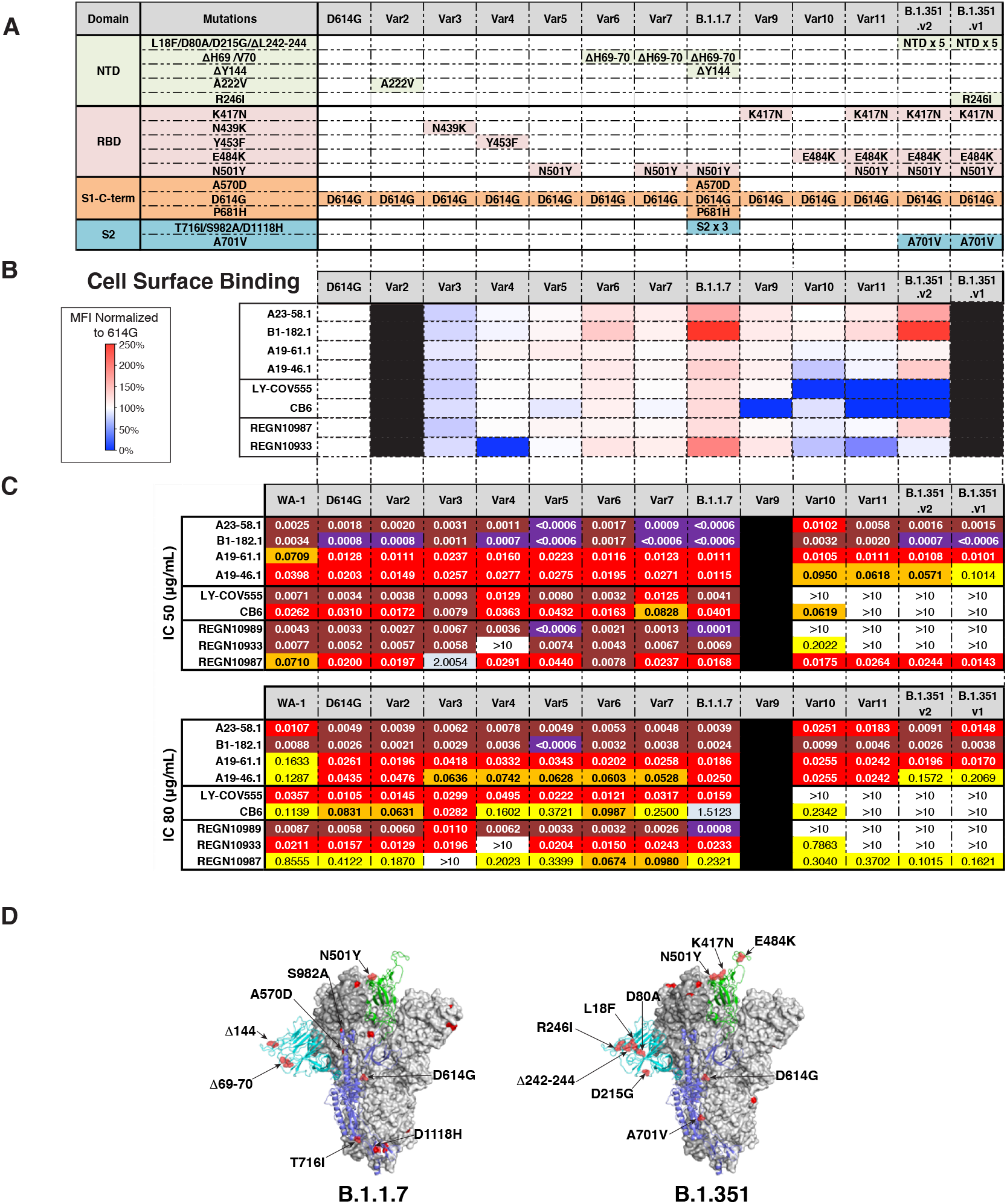
Neutralization and binding activity against spike proteins from circulating variants. **(A)** Table showing domain and mutations relative to WA-1 for each of the 13 variants tested in panels B-C. **(B)** Spike protein variants were expressed on the surface of HEK293T cells and binding to the indicated antibody was measured using flow cytometry. Data is shown as Mean Fluorescence intensity (MFI) normalized to the MFI for the same antibody against the D614G parental variant. Percent change is indicated by a color gradient from red (increased binding, Max 250%) to white (no change, 100%) to blue (no binding, 0%). Black indicates binding was not tested. **(C)** IC50 and IC80 values for the indicated antibodies against WA-1 and the 12 variants shown in (A). Ranges are indicated by colors white (>10 μg/mL), light blue (1-10 μg/mL), yellow (0.1-1 μg/mL), orange (0.05-0.1 μg/mL), red (0.01-0.05 μg/mL), maroon (0.001-0.01 μg/mL) and purple (<0.001 μg/mL). Black indicates a variant that was not tested. **(D)** Location of spike protein variant mutations on the spike glycoprotein for B.1.1.7 (left) and B.1.351 (right). P681H is not resolved in the structure and therefore its location is not noted in B.1.1.7.

Next, we assessed antibody binding to eleven cell surface expressed spike variants that have appeared subsequent to D614G (*6*–*9, 23*). We found that all control and experimental antibodies showed a minor reduction in binding (<2-fold) to Var3 (N439K/D614G). Despite this, their neutralization capacities were not significantly impacted, with the exception of REGN10987 (2.00 μg/mL) as reported previously (*24*) (Figure 2A-C, Figure S3). We also noted that while none of the experimental antibodies showed large reductions in binding, LY-CoV555, CB6 (*25*) and REGN10933 (*26*) each showed significant (>10-fold) binding deficits to one or more variants (i.e., Var4, Var9, Var10 or Var11) in these cell-based binding assays (Figure 2A,B, Figure S3).

Given the impact on antibody-spike binding, we evaluated the capacity of each antibody to neutralize virus particles pseudotyped with variant spike proteins containing one to three mutations on the D614G backbone (i.e., D614G, Var2-7 and Var10-11). Consistent with published data, REGN10933 did not neutralize Var4 (Y453F/D614G) or Var11 (K417N/E484K/N501Y/D614G) (*20, 27, 28*); CB6 did not neutralize Var11; and LY-CoV555, REGN10989, and REGN109333 showed significant potency reductions (28-fold to knockout) for neutralization of viruses containing E484K (*20, 28*) (Figure 2C). For the experimental antibodies, with the exceptions of Var5 (N501Y/D614G) and Var10 (E484K/D614G), neutralization by A23-58.1 was not significantly altered. For Var5, the IC_50_ of A23-58.1 was 5-fold lower (IC_50_ <0.0006 μg/mL) (Figure 2C). In contrast, for Var10, the IC_50_ was increased 4-fold relative to the ancestral WA-1 but was still highly potent at 0.0102 μg/mL. Neutralization by B1-182.1 maintained high-potency (IC_50_ <0.0032 μg/mL) for all variants and showed more than 4-fold improved potency for 6 of the 10 variants tested (IC_50_ <0.0008 μg/mL) (Figure 2C). Neutralization of all variants (i.e., Var2-7, Var10-11) by A19-61.1 was 3 to 6-fold more potent than WA-1 (WA-1 IC_50_ 0.0709 μg/mL; variants IC_50_ 0.0111-0.0237 μg/mL) (Figure 2A,C). Finally, neutralization by A19-46.1 was similar to WA-1 for all variants except Var10 and Var11, which were still highly potent despite having IC_50_ values that were 2 to 3-fold less active (Var10: 0.095; Var11: 0.0618; WA-1: 0.0398 μg/mL) (Figure 2C). Together, these data show the capacity of these newly identified antibodies to maintain high neutralization potency against a diverse panel of variant spike proteins.

In addition, we analyzed neutralization against two widely circulating, dominant virus variants with high-transmissibility, B.1.1.7 (a.k.a., UK VOC2020/12/01) (*6, 11*) and two versions of the B.1.351 variant (a.k.a., SA 501Y.V2), differing by the presence (B.1.351.v1) or absence (B.1.351.v2) of R246I (*7*)(Figure 2A,D). Consistent with published data, we found that LY-CoV555, CB6, REGN10989, REGN10933 and REGN10987 maintained high potency against B.1.1.7 (IC_50_ 0.0001-0.0401 μg/mL) but LY-CoV555, CB6, REGN10989 and REGN10933 were not able to neutralize either B.1.351 variant (IC_50_ >10 μg/mL) (Figure 2) (*20, 27, 28*). In comparison, A23-58.1, B1-182.1, A19-46.1 and A19-61.1 maintained similar or improved potency (IC_50_ <0.0006-0.0115 μg/mL) against B.1.1.7 relative to WA-1. A19-46.1 maintained potency against B.1.351.v2 and ∼2.5-fold lower neutralization against B.1.351.v1 (WA-1 IC_50_ 0.0398 μg/mL; B.1.351.v1 IC_50_ 0.1014 μg/mL; B.1.351.v2 IC_50_ 0.0571 μg/mL), while A23-58.1, B1-182.1 and A19-61.1 maintained high potency against both B.1.351 variants (IC_50_ <0.0006-0.0108 μg/mL) (Figure 2C). These results indicate that despite being isolated from subjects infected with early ancestral SARS-CoV-2 viruses, these antibodies have high-potency against B.1.1.7 and B.1.351 VOCs.

The two most potent antibodies, A23-58.1 and B1-182.1, shared highly similar gene family usage in their heavy and light chains, despite being from different donors (Table S1). Both use IGHV1-58 heavy chains and IGKV3-20/IGKJ1 light chains. This antibody gene family combination has previously been noted to be present in other COVID-19 convalescent subjects and has been proposed as a public clonotype (*29*–*32*). To gain structural insights on the interaction between this class of antibodies and the SARS-CoV-2 spike, we mixed A23-58.1 Fab and spike at a molar ratio of 3.6:1 and purified the complex by size-exclusion chromatography. We collected single particle cryo-EM data on a Titian Krios and determined the structure of the complex at 3.39 Å resolution (Figure 3A, Figure S4 and Table S2) and revealed that the antibody bound to spike with all RBDs in the up position, confirming the negative stain results (Figure 3A, Figure 1G). However, the cryo-EM reconstruction density of the RBD and A23-58.1 interface was poor due to conformational variation.

**Fig. 3.**
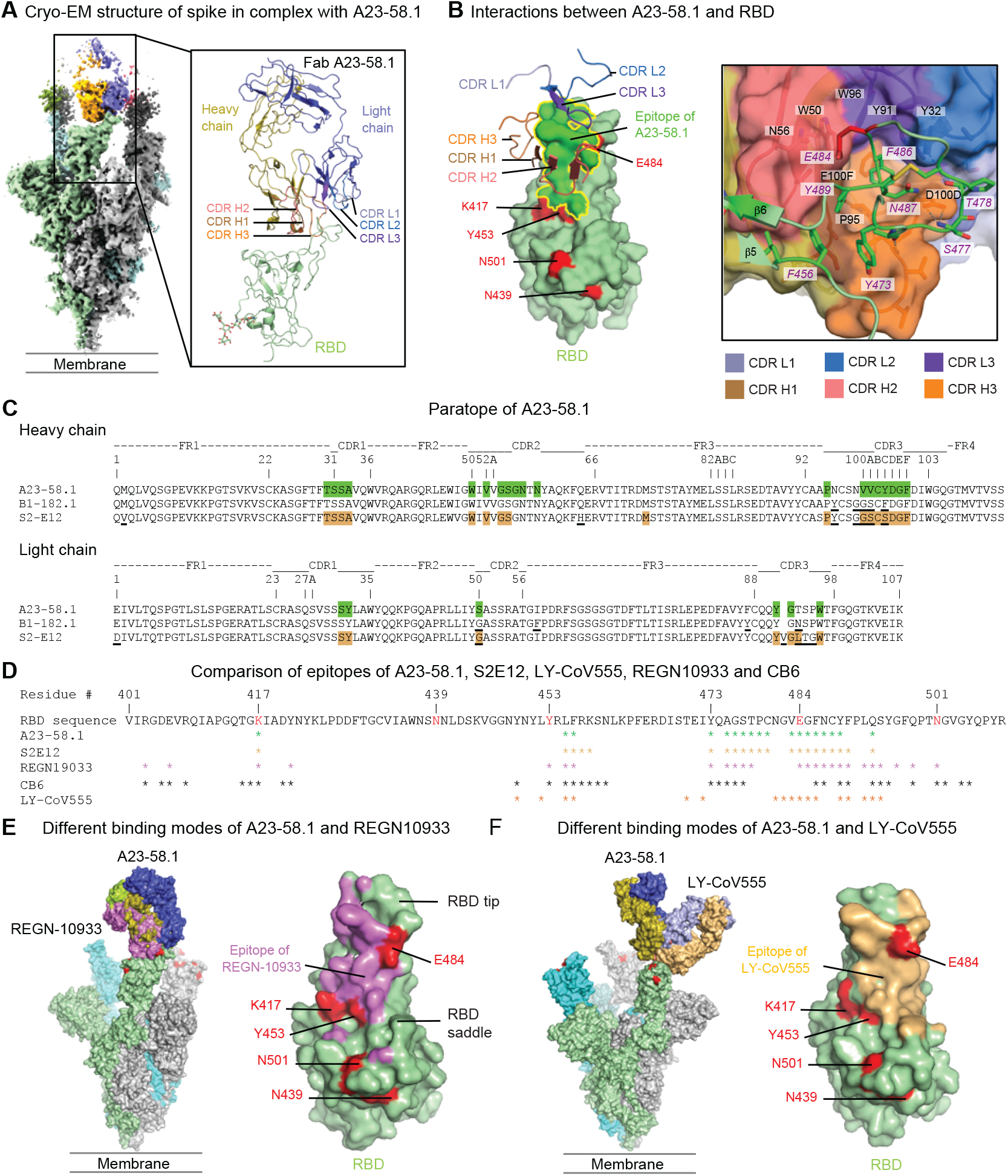
Structural basis of A23-58.1 binding. **(A)** Cryo-EM structure of A23-58.1 Fab in complex with SARS-CoV-2 HexaPro spike. Overall density map is shown to the left with protomers colored light green, gray and cyan. One of the A23-58.1 Fab bound to the RBD is shown in orange and blue. Structure of the RBD and A23-58.1 after local focused refinement was shown to the right. The heavy chain CDRs are colored brown, salmon and orange for CDR H1, CDR H2 and CDR H3, respectively. The light chain CDRs are colored marine blue, light blue and purple blue for CDR L1, CDR L2 and CDR L3, respectively. The contour level of Cryo-EM map is 5.7σ. **(B)** Interaction between A23-58.1 and RBD. All CDRs were involved in binding of RBD. Epitope of A23-58.1 is shown in bright green surface with a yellow border (left, viewing from antibody to RBD). RBD mutations in current circulating SARS-CoV-2 variants are colored red. Lys417 and Glu484 are located at the edge of the epitope. The tip of the RBD binds to a cavity formed by the CDRs (right, viewing down to the cavity). Interactions between aromatic/hydrophobic residues are prominent at the lower part of the cavity. Hydrogen bonds at the rim of the cavity are marked with dashed lines. RBD residues were labeled with italicized font. **(C)** Paratope of A23-58.1. Sequences of B1-182.1 and S2E12 were aligned with variant residues underlined. Paratope residues for A23-58.1 and S2E12 were highlighted in green and light brown, respectively. **(D)** Epitope of A23-58.1 on RBD. Epitope residues for different RDB-targeting antibodies are marked with * under the RBD sequence. **(E)** Comparison of binding modes of A23-58.1 and REGN10933. One Fab is shown to bind to the RBD on the spike. The shift of the binding site to the saddle of RBD encircled Lys417, Glu484 and Tyr453 inside the REGN10933 epitope (violet), explaining its sensitivity to the K417N, Y453F and E484K mutations. **(F)** Comparison of binding modes of A23-58.1 and LY-CoV555. One Fab is shown to bind to the RBD on the spike. Glu484 is located in the middle of LY-CoV555 epitope (light orange), explaining its sensitivity to the E484K mutation.

To resolve the antibody-antigen interface, we performed local refinement and improved the local resolution to 3.89 Å which enabled detailed analysis of the mode of antibody recognition (Figure S4). Antibody A23-58.1 bound to an epitope on the RBD that faces the 3-fold axis of the spike and is accessible only in the RBD-up conformation (Figure 3A). The interaction buried a total of 601 Å^2^ surface area from the antibody and 607 Å^2^ from the spike (Table S3). The A23-58.1 paratope constituted all six complementarity-determining regions (CDR) with both heavy chain and light chain contributing 73% and 27% of binding surface area, respectively (Figure 3B, Figure 3C and Table S3). The 14-residue-long CDR H3, which provided 48% of the heavy chain paratope, kinked at Pro95 and Phe100F (Kabat numbering scheme for antibody residues) to form a foot-like loop that is stabilized by an intra-loop disulfide bond between Cys97 and Cys100B at the arch. A glycan was observed to attach the CDR H3 Asn96 (Figure S4F). The CDRs formed an interfacial crater with a depth of ∼10 Å and a diameter of ∼20 Å at the opening. Paratope residues inside the crater were primarily aromatic or hydrophobic. With CDR H3 Pro95 and Phe100F paving the bottom, CDR H1 Ala33, CDR H2 Trp50 and Val52, and CDR H3 Val100A lined the heavy chain side of the crater (Figure 3B/C). On the light chain side, CDR L1 Tyr32 and CDR L3 residues Tyr91 and Trp96 provided 80% of the light chain binding surface (Figure 3B,C). In contrast, paratope residues at the rim of the crater are mainly hydrophilic, for example, Asp100D formed hydrogen bonds with Ser477 and Asn487 of the RBD (Figure 3B-C, Table S3).

The A23-58.1 epitope is composed of residues between β5 and β6 at the tip of RBD (Figure 3B and 3D). With the protruding Phe486 dipping into the crater formed by the CDRs, these residues formed a hook-like motif that is stabilized by an intra-loop disulfide bond between Cys480 and Cys488. Aromatic epitope residues, including Phe456, Tyr473, Phe486 and Tyr489, provided 38% of the binding surface (237.5 Å^2^) (Figure 3B and 3D, Table S3). Lys417 and Glu484, which are located at the outer edge of the epitope, contributed only 3.7% of the binding surface (Figure 3B and Table S3). Overall, the cryo-EM analysis provided structural basis for the potent neutralization of the E484K mutant by A23-58.1. The binding mode of A23-58.1 is very similar to that of a previously reported IGHV1-58/IGKV3-20-derived antibody, S2E12 (*29*) confirming that they are members of the same structural class (Figure 3C and 3D). In addition, sequence analysis indicates that B1-182.1 is likely also a member of this class – and thus shares the same mode of recognition. In fact, B1-182.1 share a nearly identical IGHD2-15-derived CDR H3 sequence with S2E12 (Figure 3C). The lack of impact of emerging resistance mutations on B1-182.1 can be explained by the same mechanism whereby A23-58.1 antibody also is not impacted by these variants. Interestingly, while A23-58.1 and B1-182.1 both have 10-residue-long CDR L3s, S2E12 includes the insertion of Val91A (Figure 3C) and the longer CDR L3 may therefore lead to differences in binding/neutralization capacity for viruses with mutations at the Glu484 location where CDR L3 contacts.

We next sought to understand the structural mechanisms by which A23-58.1 overcomes mutations that cause reduced antibody potency against virus variants. To do this, we superposed the antibody-RBD complex structures of CB6 (PDB ID 7C01) (*25*), REGN10933 (PDB ID 6XDG) (*26, 27*) and LY-CoV555 (PDB ID 7KMG) (*18*) with the A23-58.1 structure over the RDB region. Both REGN10933 and CB6 bind to the same side of the RBD that A23-58.1 contacted (Figure 3D and 3E). However, their binding surfaces were all shifted towards the saddle of the open RBD and encircled residues Lys417, Tyr453, Glu484 and Asn501 within the epitope (Figure 3D and 3E), mutations K417N and Y453F potently abolished key interactions and led to the loss of neutralization for both REGN10933 and CB6 (Figure 2). In contrast, LY-CoV555 approached the RBD from a different angle with its epitope centered around Glu484 (Figure 3D and 3F). Modeling indicated that mutation E484K may abolish key interactions with Arg50 and Arg96 of LY-CoV555 and cause a clash with CDR H3 of LY-CoV555. These structural data suggest that the unique binding modes of A23-58.1 and potentially B1-182.1 derived from the same germline enabled their high effectiveness against the new SARS-CoV-2 variants.

We next used the structural analysis to investigate the relative contribution of predicted contact residues on binding and neutralization (Figure 3D). Cell surface expressed spike binding by A23-58.1 and B1-182.1 were knocked out by F486R, N487R, and Y489R (Figure 4A, Figure S5), resulting in a lack of neutralization for viruses pseudotyped with spikes containing these mutations (Figure 4B). In contrast, binding and neutralization of A19-46.1 and A19-61.1 were minimally impacted by these changes (Figure 4A,B, Figure S5). CB6, LY-CoV555 and REGN10933 binding and neutralization were also impacted by the three mutations, consistent with the structural analysis that these residues are commonly shared contact(s) among the impacted antibodies. Taken together, the shared binding and neutralization defect imposed by these mutations on A23-58.1 and B1-182.1 suggests that the hook-like motif and CDR crater are critical for the binding of antibodies within the VH1-58 public class.

**Fig. 4.**
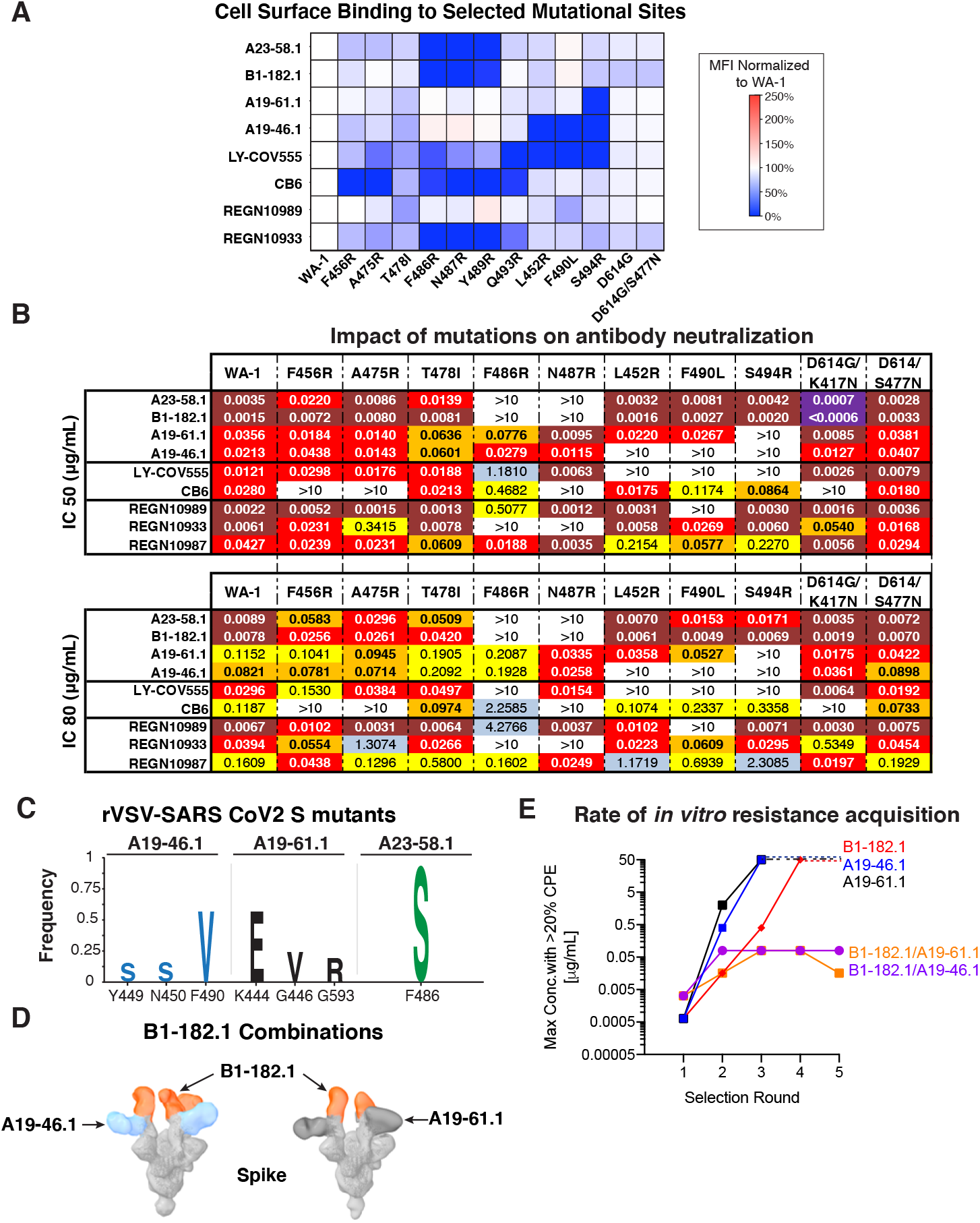
Critical binding residue determination and mitigation of escape risk using dual antibody combinations. **(A)** The indicated Spike protein mutations predicted by structural analysis were expressed on the surface of HEK293T cells and binding to the indicated antibody was measured using flow cytometry. Data is shown as Mean Fluorescence intensity (MFI) normalized to the MFI for the same antibody against the WA-1 parental binding. Percent change is indicated by a color gradient from red (increased binding, Max 250%) to white (no change, 100%) to blue (no binding, 0%). **(B)** IC_50_ and IC_80_ values for the indicated antibodies against WA-1 and the 10 mutations. Ranges are indicated by colors white (>10 μg/mL), light blue (1-10 μg/mL), yellow (0.1-1 μg/mL), orange (0.05-0.1 μg/mL), red (0.01-0.05 μg/mL), maroon (0.001-0.01 μg/mL) and purple (<0.001 μg/mL). **(C)** Replication competent vesicular stomatitis virus (rcVSV) whose genome expressed SARS-CoV-2 WA-1 was incubated with serial dilutions of the indicated antibodies and wells with cytopathic effect (CPE) were passaged forward into subsequent rounds (Figure S6) after 48-72 hours. Total supernatant RNA was harvested and viral genomes shotgun sequenced to determine the frequency of amino acid changes. Shown are the spike protein amino acid/position change and frequency as a logo plot. **(D)** Negative stain 3D reconstruction of the ternary complex of spike with Fab B1-182.1 and A19-46.1 (left) or A19-61.1 (right). **(E)** rcVSV SARS-CoV-2 was incubated with increasing concentrations (1.3e-4 to 50 μg/mL) of either single antibodies (A19-46.1, A19-61.1 and B1-182.1) and combinations of antibodies (B1-182.1/A19-46.1 and B1-182.1/A19-61.1). Every 3 days, wells were assessed for CPE and the highest concentration well with the >20% CPE was passaged forward onto fresh cells and antibody containing media. Shown is the maximum concentration with >20% CPE for each of the test conditions in each round of selection. Once 50 μg/mL has been reached, virus was no longer passaged forward and a dashed line is used to indicate maximum antibody concentration was reached in subsequent rounds.

Next, based on structural modeling of the negative stain EM density (Figure 1G), we chose several mutants to investigate the determinants of binding for A19-46.1 and A19-61.1. Under conditions where A23-58.1 and B1-182.1 were not impacted, we found that L452R, F490R and S494R knocked out binding for A19-46.1 and S494R knocked out binding for A19-61.1 (Figure 4A, Figure S5). In addition, the partial overlap of susceptibility to the selected mutations between A19-46.1 and A19-61.1 is in agreement with the antibody competition data showing similar, but distinct profiles (Figure 1F) and indicates that these antibodies represent distinct antibody classes.

To explore resistance mechanisms that might be generated during the course of infection, we applied antibody selection pressure to replication competent vesicular stomatitis virus (rcVSV) expressing the WA-1 SARS-CoV-2 spike (rcVSV-SARS2) (*33*) to identify spike mutations that confer *in vitro* resistance against A23-58.1, B1-182.1, A19-46.1 or A19-61.1 (Figure S6). rcVSV-SARS2 was incubated with increasing concentrations of antibody, and cultures from the highest concentration of antibody with >20% cytopathic effect (CPE) were carried forward into a second round of selection to drive resistance (*27*)(Figure S6). A shift to higher antibody concentrations required for neutralization indicated the presence of resistant viruses. To define and determine the relative frequency of mutations that accumulated at positions within the spike of resistant viruses, we performed Illumina-based shotgun sequencing (Figure S6). Variants present at a frequency of greater than 10% and increasing from round 1 to round 2 were considered to be positively selected resistant viruses. For A19-46.1, three selection mutations were generated: Y449S (freq. 15%), N450S (freq. 16%) and F490V (freq. 58%) (Figure 4C, Figure S7). The most dominant, F490V, was found in 58% of sequences and is consistent with the previous finding that F490R knocked out binding and neutralization of A19-46.1 (Figure 4A,C). These residues are clustered near one another on RBD and would be expected to be accessible when RBD is in the up or down position (Figure S7). Several of these contacts are shared by Class II RBD antibodies (*22, 34*) and REGN10933 (*26, 35*). Despite these shared contacts, A19-46.1 is able to neutralize variants that REGN10933 cannot (i.e., Var10, Var11 and B.1.351) (Figure 2A-C), indicating that A19-46.1 makes critical contacts that are able to overcome resistance mutations that affect REGN10933.

Three residues were positively selected in the presence of A19-61.1: K444E (freq. 57%), G446V (freq. 24%) and G593R (freq. 19%) (Figure 4A) and did not overlap with those selected by A19-46.1. G593R is located outside the RBD domain and the others are clustered nearby S494R, identified previously (Figure 4A, Figure S7). The highest frequency change was at K444 and represented 57% of the sequences. This residue has been shown to be critical for the binding of Class III RBD antibodies such as REGN10987 (*22, 26, 27, 35*). Taken together with the ability of A19-61.1 to block ACE2 binding (Figure 1F, Figure S2) and differential neutralization between the Class III antibody REGN10987 and A19-61.1 against Var3 (N439K/D614G) (i.e., significantly reduced neutralization with REGN10987) (Figure 2C), A19-61.1 likely targets an epitope distinct epitope from REGN10987 and other Class III RBD antibodies.

Finally, a single mutation, F486S (freq. 91%) was positively selected for when virus was incubated in the presence of A23-58.1. This is in agreement with our structural analysis (Figure 3B) that showed that F486 is located at the tip of RBD “hook” and contributes to the binding interface in the antibody “crater”. This is dominated by aromatic side chains that form the “hook” and “crater” interface (Figure 3A,B). Therefore, one possible explanation for the loss in activity is through the replacement of a hydrophobic aromatic residue (i.e., phenylalanine) with a small polar side chain (i.e., serine) (Figure 3C).

To probe the relevance of in vitro derived resistance variants to the potential for clinical resistance we next examined spikes derived from circulating virus variants. We investigated the relative frequency of variants containing the mutations present in the GISAID sequence database using the COVID-19 Viral Genome Analysis Pipeline (cov.lanl.gov)(*23*) in which, as of February 11, 2021, there were 417,702 entries. Out of these, the original WA-1 residues critical for A19-46.1 activity (i.e., Y449, N450, L452, F490 and S494) were present in 99.81-99.99% of sequences available. Of the residues identified in our experiments to mediate resistance to A19-46.1, Y449S, N450S, L452R, F490L/V and S494R, only F490V has been noted in the database (5 sequences, 0.001%) (Figure 4, Figure S7). For A19-61.1, the ancestral WA-1 residues, K444, G446, S494 and G593, were present in 99.81-100% of entries. Of the resistance-inducing residues identified, i.e., K444E, G446V, S494R, and G593R, only G446V has been noted in the database (106 sequences, 0.03%) (Figure 4, Figure S7). Finally, for A23-58.1 and B1-182.1 the ancestral WA-1 residues F486, N487 and Y489 were present in >99.99% of sequences and none of the binding/resistance mutations identified in our experiments were noted in the database. The relative lack of resistance mutations found in circulating viruses suggests that the *in vitro* derived mutations exact a fitness cost on the virus and are not tolerated during infection but could also reflect either under-sampling or the absence of other sources of selection pressure.

Viral genome sequencing has suggested the possibility that in addition to spread via transmission, convergent selection of *de novo* mutations may be occurring(*6*–*9, 13, 23, 36*). Therefore, effective therapeutic antibody approaches might require new antibodies or combinations of antibodies to mitigate the impact of mutations. Based on their complementary modes of spike recognition and breadth of neutralizing activity, we hypothesized that combination of B1-182.1 with either A19-46.1 or A19-61.1 would decrease the rate of *in vitro* resistance acquisition compared to each antibody alone. As a first test, we used negative stain EM 3D reconstructions to determine whether these combinations are able to bind simultaneously to spike protein. Consistent with the competition data (Figure 1F), we found that the Fabs in both combinations were able to engage spike simultaneously with RBD in the up position (Figure 4D). Furthermore, we noted that binding was in a 3:1 Fab:spike ratio in most of the observed particles (Figure 4D), revealing that the epitopes of A19-46.1 and A19-61.1 on the spike are accessible in both RBD up and down positions (Figure 1G and Figure 4D). This suggests that the combination allows alternative preferential mode of RBD engagement (i.e., RBD up vs. RBD down) by A19-46.1 and A19-61.1 that is not seen in the absence of B1-182.1 or A23-58.1.

Next, we determined the rates of *in vitro* resistance acquisition of combined treatments compared to individual antibodies using an rcVSV SARS-CoV-2 resistance generation approach. We evaluated the capacity of individual antibodies or combinations to prevent the appearance of rcVSV SARS –CoV-2-induced cytopathic effect (CPE) throughout multiple rounds of passaging in the presence of increasing concentrations of antibodies. In each round, the well with the highest concentration of antibody with at least 20% CPE was carried forward into the next round. We found that wells with A19-61.1 or A19-46.1 single antibody treatment reached the 20% CPE threshold in their 50 μg/mL well after 3 rounds of selection (Figure 4E). Similarly, B1-182.1 single antibody treatment reached >20% CPE in the 50 μg/mL wells after 4 rounds (Figure 4E). Conversely, for both dual treatments (i.e., B1-182.1/A19-46.1 or B1-182.1/A19-61.1) the 20% CPE threshold was reached only at a concentration of 0.08 μg/mL and did not progress to higher concentrations despite 5 rounds of passaging (Figure 4E). While further data are required, these results suggest that such combinations may lower the risk that a natural variant will lead to the complete loss of neutralizing activity and suggests a path forward for these antibodies as combination therapies.

Worldwide genomic sequencing has revealed the occurrence of SARS-CoV-2 variants that increase transmissibility and reduce potency of vaccine-induced and therapeutic antibodies (*10*– *16*). Recently, there has been a significant concern that antibody responses to natural infection and vaccinations using ancestral spike sequences may have focused responses that are overcome by mutations present in more recent isolates (e.g., E484K in B.1.351) (*12*–*16*). As a first step to address the risk of reduced antibody potency against new variants, we isolated and defined new antibodies with neutralization breadth covering newly emerging SARS-CoV-2 variants. Increased potency and breadth were mediated by binding to regions of the RBD tip that are offset from E484K, which is a major determinant of resistance in VOCs (*12*–*16*). Our results show that highly potent neutralizing antibodies with activity against these variants was present in at least 3 of four convalescent subjects who had been infected with ancestral variants of SARS-CoV-2. Furthermore, two antibodies from different subjects used VH genes associated with previously described public clonotypes (*29, 30*). Overall, these data establish the rationale for a vaccine boosting regimen that may be used to selectively induce immune responses that increase the breadth and potency of antibodies targeting the RBD region of the spike glycoprotein. Furthermore, since both variant sequence analysis and *in vitro* time to escape experiments suggest that combinations of these antibodies may have a lower risk for loss of neutralizing activity, these antibodies represent a potential means to achieve both breadth against current VOCs and to mitigate risk against those that may develop in the future.

## Acknowledgments

We would like to thank the staff of the Clinical Trials Program of the Vaccine Research Center and the volunteers that made this research possible. We also appreciate the assistance of Dr. Ruth Hunegnaw for assistance with figure preparation. We are grateful to Tara L. Fox of NCEF for collecting cryo-EM data and for technical assistance with cryo-EM data processing. We would like to thank Avan Antia, Rachel L. Davis and Farida Laboune for technical assistance with sequencing.

## Author contributions

T.R., E.P., A.D., L.N., N.D.R., R.M., J.M., C.S., L.W., K.C. and E.C. designed and performed cell sorting experiments. A.R., S.D. and C.S. performed and analyzed sequencing data. Proteins, antibody and other reagents were produced by W.S., I.T., L.W., T.Z., A.O., E.P., T.R., J.M., O.A., L.C., A.D., E.S.Y. Y.Z., B.Z., A.N. and T.L., J.M., L.W., T.Z., Y.Z., W.S., E.S.Y., A.P., O.O., A.R., C.S., S.D., S.N., C.H., D.M., C.T., C.S. and D.W. conceived of, designed experiments, performed experiments, data analysis and reporting. M.G., A.W., L.N. and I.G. for research subject recruitment, collection of samples and maintenance of the sample repository. T.Z. and Y.T. led negative stain electron microscopy and cryo-EM studies. J.M., N.J.S., J.R.M., D.D., B.S.G, A.M., P.K. and R.S.B. supervised experiments. J.M., N.J.S., T.Z., L.W. and C.S. wrote the manuscript with help from all authors.

## Data and materials availability

All data is available in the main text or the supplementary materials. Atomic coordinates and cryo-EM maps of the reported structure have been deposited into the Protein Data Bank and Electron Microscopy Data Bank under the session codes PDB 7LRT and EMD-23499 for SARS-CoV-2 spike in complex with antibody A23-58.1, and PDB 7LRS and EMD-23498 for local refinement of the RDB-antibody A23-58.1 region. Plasmids are available from N.J.S. under a materials transfer agreement with the National Institutes of Health.

## Supplementary Materials

Materials and Methods

Figures S1-S7

Tables S1-S3

References (36-52)

