## Supplemental Materials and Methods for "Antibodies with potent and broad neutralizing activity against antigenically diverse and highly transmissible SARS-CoV-2 variants"

†Equal contributions

**This PDF file includes:**

Materials and Methods

Figs. S1 to S7

Tables S1 to S3

Captions for Data S1 to S6 and Table S1 to S3

#### Materials and Methods

Isolation of PBMCs from SARS CoV-2 subjects

Human convalescent sera samples were obtained 30 to 50 days following symptom onset from adults with previous mild to moderate SARS-CoV-2 infection. Specimens were collected after subjects provided written informed consent under institutional review board approved protocols at the National Institutes of Health Clinical Center (NCT00067054) and University of Washington (Seattle) (Hospitalized or Ambulatory Adults with Respiratory Viral Infection [HAARVI] study). Whole blood was collected in vacutainer tubes, which were inverted gently to remix cells prior to standard Ficoll-Hypaque density gradient centrifugation (Pharmacia; Uppsala, Sweden) to isolate PBMCs. PBMCs were frozen in heat-inactivated fetal calf serum containing 10% dimethylsulfoxide in a Forma CryoMed cell freezer (Marietta, OH). Cells were stored at -≤140°C

Expression and Purification of Protein

For expression of soluble SARS CoV-2 S-2P protein, manufacturer's instructions were followed. Briefly, plasmid was transfected using Expifectamine into Expi293 cells (Life Technology) and the cultures enhanced 16-24 hours post-transfection. Following 4-5 days incubations at 120 rpm, 37 °C, 9% CO2, supernatant was harvested, clarified via centrifugation, and buffer exchanged into 1X PBS. Protein of interests were then isolated by affinity chromatography using Streptactin resin (Life science) followed by size exclusion chromatography on a Superose 6 increase 10/300 column (GE healthcare).

Expression and purification of biotinylated S-2P, NTD, RBD-SD1 and Hexapro used in binding assays were produced by an in-column biotinylation method as previously described (5).

Using full-length SARS-Cov2 S and human ACE2 cDNA ORF clone vector (Sino Biological, Inc) as the template to generate S1 or ACE2 dimer proteins. The S1 PCR fragment (1~681aa) was digested with Xbal and BamHI and cloned into the VRC8400 with HRV3C-his (6X) or Avi-HRV3C-his(6X) tag on the C-terminal. The ACE2 PCR fragment (1~740aa) was digested with Xbal and BamHI and cloned into the VRC8400 with Avi-HRV3C-single chain-human Fc-his (6x) tag on the C-terminal. All constructs were confirmed by sequencing. Proteins were expressed in Expi293 cells by transfection with expression vectors encoding corresponding genes. The transfected cells were cultured in shaker incubator at 120 rpm, 37 °C, 9% $CO_2$ for 4~5 days. Culture supernatants were harvested and filtered, and proteins were purified through a Hispur Ni-NTA resin (Thermo Scientific) and following a Hiload 16/600 Superdex 200 column (GE healthcare, Piscataway NJ) according to manufacturer's instructions. The protein purity was confirmed by SDS-PAGE.

Probe conjugation

SARS CoV-2 Spike trimer (S-2P) and subdomains (NTD, RBD-SD1, S1) were produced by transient transfection of 293 Freestyle cells as previously described (2). Avi-tagged S1 was biotinylated using the BirA biotin-protein ligase reaction kit (Avidity) according to the manufacturer's instructions. The S-2P, RBD-SD1, and NTD proteins were produced by an in-column biotinylation method as previously described (5). Successful biotinylation was confirmed using Bio-Layer Interferometry, by testing the ability of biotinylated protein to bind to streptavidin sensors. Retention of antigenicity was confirmed by testing biotinylated proteins against a panel of cross-reactive SARS-CoV and SARS CoV-2 human monoclonal antibodies. Biotinylated probes were conjugated using either allophycocyanin (APC)-, Ax647, BV421-, BV786, BV711-, or BV570-labeled streptavidin. Reactions were prepared at a 4:1 molecular ratio of biotinylated protein to streptavidin, with every monomer labeled. Labeled streptavidin was added in ⅕ increments. The streptavidin-protein mix was incubated at in the dark at 4°C (rotating) for 20 minutes in between each addition. Optimal titers were determined using splenocytes from immunized mice and validated with SARS CoV-2 convalescent human PBMC.

Isolation of and sequencing of antibodies by single B cell sorting

Cryopreserved human PBMCs from four COVID-19 convalescent donors Subjects 1-4 were thawed and stained with Live/DEAD Fixable Aqua Dead Cell Stain kit (Cat# L34957, ThermoFisher). After washing, cells were stained with a cocktail of anti-human antibodies, including CD3 (cat # 317332, Biolegend), CD8 (cat # 301048, Biolegend), CD56 (cat # 318340, Biolegend), CD14 (cat # 301842, Biolegend), CD19 (Cat# IM2708U, Beckman Coulter), CD20 (cat # 302314, Biolegend), IgG (Cat# 555786, BD Biosciences), IgA (Cat# 130-114-001, Miltenyi), IgM (Cat# 561285, BD Biosciences) and subsequently stained with fluorescently labeled SARS-CoV-2 S-2P (APC or Ax647), S1 (BV786 or BV570), RBD-SD1 (BV421) and NTD (BV711 or BV421) probes. Antigen-specific memory B cells (CD3-CD19+CD20+ IgG+/IgA+ and S-2P+ and/or RBD+ for the donors Subjects 1-3, S-2P+ and/or NTD+ for the donor Subject 4) were sorted using a FACSymphony S6 (BD Sciences) into Buffer TCL (Qiagen) with 1% 2-mercaptoethanol (ThermoFisher Scientific). Nucleic acids were purified using RNAClean magnetic beads (Beckman Coulter) followed by reverse transcription using oligo-dT linked to a custom adapter sequence and template switching using SMARTScribe RT (Takara). PCR amplification was carried out using SeqAmp DNA Polymerase (Takara). A portion of the amplified cDNA was enriched for B cell receptor sequences using forward primers complementary to the template switch oligo and reverse primers against the IgA (GAGGCTCAGCGGGAAGACCTTGGGGCTGGTCGG) IgG, Igκ, and Igλ (38) constant regions. Enriched products were made into Illumina-ready sequencing libraries using the Nextera XT DNA Library Kit with Unique Dual Indexes (Illumina). The Illumina-ready libraries were sequenced by paired end 150 cycle MiSeq reads. The resulting reads were demultiplexed using an in-house script and V(D)J sequences were assembled using BALDR in unfiltered mode (39). Poor or incomplete assemblies or those with low read support were removed, and the filtered contigs were re-annotated with SONAR v4.2 in single cell mode (40). A subset of the final antibodies were manually selected for synthesis based on multiple considerations, including gene usage, somatic hypermutation levels, CDRH3 length, convergent rearrangements, and specificity implied by flow cytometry.

Synthesis, cloning and expression of monoclonal antibodies

Variable heavy chain sequences were human codon optimized, synthesized and cloned into a VRC8400 (CMV/R expression vector)-based IgG1 vector containing an HRV3C protease site (41) as previously described (37). Similarly, variable lambda and kappa light chain sequences were human codon optimized, synthesized and cloned into CMV/R-based lambda or kappa chain expression vectors, as appropriate (Genscript). Previously published antibody vectors for LY-CoV555(18) and mAb114 (37) were used. The antibodies: REGN10989, REGN10987 and REGN10933, were produced from published sequences (26) and kindly provided by Devin Sok from Scripps. For antibodies where vectors were unavailable (e.g., S309, CB6), published amino acids sequences were used for synthesis and cloning into corresponding pVRC8400 vectors (21,25). For antibody expression, equal amounts of heavy and light chain plasmid DNA were transfected into Expi293 cells (Life Technology) by using Expi293 transfection reagent (Life Technology). The transfected cells were cultured in shaker incubator at 120 rpm, 37 °C, 9% $CO_2$ for 4~5 days. Culture supernatants were harvested and filtered, mAbs were purified over Protein A (GE Health Science) columns. Each antibody was eluted with IgG elution buffer (Pierce) and immediately neutralized with one tenth volume of 1M Tris-HCL pH 8.0. The antibodies were then buffer exchanged as least twice in PBS by dialysis.

Assignment of major binding determinant using MSD binding assay

MSD 384-well streptavidin-coated plates (MSD, cat# L21SA) were blocked with MSD 5% Blocker A solution (MSD, cat# R93AA), using 35 ul per well. These plates were then incubated for 30 to 60 minutes at room temperature. Plates were washed with 1x Phosphate Buffered Saline + 0.05% Tween 20 (PBST) on a Biotek 405TS automated microplate washer. Five SARS CoV-2 capture antigens were used. Capture antigens consisted of VRC-produced S1, S-2P, S6P (Hexapro), RBD, and NTD. All antigens were AVI-tag biotinylated using BirA (Avidity, cat # BirA500) AVI-tag specific biotinylation following manufacturer's instructions except S1. For S1, an Invitrogen FluoReporter™ Mini-Biotin-XX Protein Labeling Kit (Thermo Fisher, cat # F6347) was utilized to achieve random biotinylation. Antigen coating solutions were prepared for S1, S-2P, S6P, RBD, and NTD at optimized concentrations of 0.5, 0.25, 1, 0.5, and 0.25 ug/mL, respectively. These solutions were then added to MSD 384-well plates, using 10 µL per well. Each full antigen set is intended to test one plate of experimental SARS CoV-2 monoclonal antibodies (mAbs) at one dilution. Once capture antigen solutions were added, plates were incubated for 1 hour at room temperature on a Heidolph Titramax 1000 (Heidolph, part # 544-12200-00) vibrational plate shaker at 1000 rpm. During this time, experimental SARS CoV-2 mAb dilution plates were prepared. Using this initial plate, 3 dilution plates were created at dilution factors of 1:100, 1:1000, and 1:10000. Dilutions were performed in 1% assay diluent (MSD 5% Blocker A solution diluted 1:5 in PBST). Positive control mAbs S652-109 (SARS Cov-2 RDB specific) and S652-112 (SARS CoV-2 S1, S-2P, S6P, and NTD specific) and negative control mAb VRC01 (anti-HIV) were added to all dilution plates at a uniform concentration of 0.05 µg/mL. Once mAb dilution plates were prepared, MSD 384-well plates were washed as above. The content of each 96-well dilution plate was added to the MSD 384-well plates, using 10 µL per well. MSD 384-well plates were then incubated for 1 hour at room temperature on vibrational plate shaker at 1000 rpm. MSD 384-well plates were washed as above, and MSD Sulfo-Tag labeled goat anti-human secondary detection antibody (MSD, cat# R32AJ) solution was added to plates at a concentration of 0.5 ug/mL, using 10 µL per well. Plates were again incubated for 1 hour at room temperature on vibrational plate shaker at 1000 rpm. MSD 1x Read Buffer T (MSD, cat# R92TC) was added to MSD 384-well plates, using 35 µL per well. MSD 384-well plates were then read using MSD Sector S 600 imager. Gross binding epitope of S-2P or Hexapro positive antibodies was assigned into the following groups: RBD (i.e., RBD+ or RBD+/S1+ AND NTD-), NTD (i.e., NTD+ or NTD+/S1+ AND RBD-), S2 (i.e., S1-, RBD- AND NTD-) or indeterminant (i.e., mixed positive). Antibodies lacking binding to any of the antigens were assigned to the "no binding" group.

Full-length S constructs

cDNAs encoding full-length S from SARS CoV-2 (GenBank ID: QHD43416.1) were synthesized, cloned into the mammalian expression vector VRC8400 (42,43) and confirmed by sequencing. S containing D614G amino acid change was generated using the wt S sequence. Other variants containing single or multiple aa changes in the S gene from the S wt or D614G were made by mutagenesis using QuickChange lightning Multi Site-Directed Mutagenesis Kit (cat # 210515, Agilent). The S variants, N439K, Y453F, A222V, E484K, K417N, S477N, N501Y, delH69/V70, N501Y-delH69/V70, N501Y-E484K-K417N, B.1.1.7 (H69del-V70del-Y144del-N501Y-A570D-P681H-T716I-S982A-D1118H), B.1.351.v1 (L18F-D80A-D215G-(L242-244)del-R246I-K417N-E484K-N501Y-A701V) and B.1.351.v2 (L18F-D80A-D215G-(L242-244)del-K417N-E484K-N501Y-A701V) were generated based on S D614G while the variants, F456R, A475R, T478I, F486R, N487R, L452R, F490L, S494R on S wt. These full-length S plasmids were used for pseudovirus production and for cell surface binding assays.

Pseudovirus neutralization assay

S-containing lentiviral pseudovirions were produced by co-transfection of packaging plasmid pCMVdR8.2, transducing plasmid pHR' CMV-Luc, a TMPRSS2 plasmid and S plasmids from SARS CoV-2 variants into 293T cells using Fugene 6 transfection reagent (Promega, Madison, WI) (44-46). 293T-ACE2 cells, provided by Dr. Michael Farzan, were plated into 96-well white/black Isoplates (PerkinElmer, Waltham, MA) at 5,000 cells per well the day before infection of SARS CoV-2 pseudovirus. Serial dilutions of mAbs were mixed with titrated pseudovirus, incubated for 45 minutes at 37°C and added to 293T-ACE2 cells in triplicate. Following 2 h of incubation, wells were replenished with 150 ml of fresh media. Cells were lysed 72 h later, and luciferase activity was measured with Microbeta (Perking Elmer). Percent neutralization and neutralization IC50s, IC80s were calculated using GraphPad Prism 8.0.2.

Cell surface binding

HEK293T cells were transiently transfected with plasmids encoding full length SARS CoV-2 spike variants using lipofectamine 3000 (L3000-001, ThermoFisher) following manufacturer's protocol. After 40 hours, the cells were harvested and incubated with monoclonal antibodies (1 μg/ml) for 30 minutes. After incubation with the antibodies, the cells were washed and incubated with an allophycocyanin conjugated anti-human IgG (709-136-149, Jackson Immunoresearch Laboratories) for another 30 minutes. The cells were then washed and fixed with 1% paraformaldehyde (15712-S, Electron Microscopy Sciences). The samples were then acquired in a BD LSRFortessa X-50 flow cytometer (BD biosciences) and analyzed using Flowjo (BD biosciences). Mean fluorescent intensity (MFI) for antibody binding to S wt or D614G was set up as 100%. The MFI of the antibody binding to each variant was normalized to S wt or D614G.

<u>Inhibition of S protein binding to cell surface ACE2</u>

Serial dilutions of mAb IgG and Fab were mixed with pre-titrated biotinylated S trimer (S-2P), incubated for 30 minutes at RT and added to BHK21 cells stably expressing hACE2 on cell surface. Following 30 minutes of incubation on ice, the cells were washed and incubated with an BV421 conjugated Streptavidin (cat # 563259, BD Biosciences) for another 30 minutes. The cells were then washed and fixed with 1% paraformaldehyde (15712-S, Electron Microscopy Sciences). The samples were then acquired in a BD LSRFortessa X-50 flow cytometer (BD biosciences) and analyzed using Flowjo (BD biosciences). Mean fluorescent intensity (MFI) for S protein binding to cell surface was set up as 100%. Percent inhibition of S protein binding to cell surface ACE2 by mAb IgG and EC50s were calculated using GraphPad Prism 8.0.2.

<u>Live virus neutralization assay</u>

Full-length SARS CoV-2 virus based on the Seattle Washington strain was designed to express nanoluciferase (nLuc) and was recovered via reverse genetics and described previously (17). Virus titers were measured in Vero E6 USAMRIID cells, as defined by plaque forming units (PFU) per ml, in a 6-well plate format in quadruplicate biological replicates for accuracy. For the 96-well neutralization assay, Vero E6 USAMRID cells were plated at 20,000 cells per well the day prior in clear bottom black walled plates. Cells were inspected to ensure confluency on the day of assay. Serially diluted mAbs were mixed in equal volume with diluted virus. Antibody-virus and virus only mixtures were then incubated at 37°C with 5% $CO_2$ for one hour. Following incubation, serially diluted mAbs and virus only controls were added in duplicate to the cells at 75 PFU at 37°C with 5% $CO_2$. After 24 hours, cells were lysed, and luciferase activity was measured via Nano-Glo Luciferase Assay System (Promega) according to the manufacturer specifications. Luminescence was measured by a Spectramax M3 plate reader (Molecular Devices, San Jose, CA). Virus neutralization titers were defined as the sample dilution at which a 50% reduction in RLU was observed relative to the average of the virus control wells.
Live virus neutralization assays described above were performed with approved standard operating procedures for SARS CoV-2 in a biosafety level 3 (BSL-3) facility conforming to requirements recommended in the Microbiological and Biomedical Laboratories, by the U.S. Department of Health and Human Service, the U.S. Public Health Service, and the U.S. Center for Disease Control and Prevention (CDC), and the National Institutes of Health (NIH).

Production of Fab fragments from monoclonal antibodies

To generate mAb-Fab, IgG was incubated with HRV3C protease (EMD Millipore) at a ratio of 100 units per 10 mg IgG with HRV 3C Protease Cleavage Buffer (150mM NaCl, 50mM

Tris-HCl, pH 7.5) at 4°C overnight. Fab was purified by collecting flowthrough from Protein A column (GE Health Science), and Fab purity was confirmed by SDS-PAGE.

Competitive binding assay using biolayer interferometry

Antibody cross-competition was determined based on biolayer interferometry using a fortéBio Octet HTX instrument. His1K biosensors (fortéBio) were equilibrated for >600 s in Blocking Buffer (1% BSA (Sigma) + 0.01% Tween-20 (Sigma) + 0.01% Sodium Azide (Sigma) + PBS (Gibco), pH7.4) prior to loading with his tagged S-2P protein (10 μg/mL in Blocking Buffer) for 1200s. Following loading, sensors were incubated for 420s in Blocking Buffer prior to incubation with competitor mAbs (30 mg/mL in Blocking Buffer) for 1200s. Sensors were then incubated in Blocking buffer for 30s prior to incubation with the analyte mAbs (30 mg/mL in Blocking Buffer) for 1200s. Percent competition (PC) of analyte mAbs binding to competitor-bound S-2P was determined using the equation: PC = 100 − [(analyte mAb binding in the presence competitor mAb) ∕ (analyte mAb binding in the absence of competitor mAb)] × 100. All the assays were performed in duplicate and with agitation set to 1,000 rpm at 30°C

Determination Fab fragment kinetics of binding

A fortéBio Octet HTX instrument was used to measure binding kinetics of the Fab of A23-58.1, B1-182.1, A19-46.1 and A19-61.1 to SARS CoV-2 S-2P protein. SA biosensors (fortéBio) were equilibrated for >600 s in Blocking Buffer (1% BSA (Sigma) + 0.01% Tween-20 (Sigma) + 0.01% Sodium Azide (Sigma) + PBS (Gibco), pH7.4) prior to loading with biotinylated S-2P protein (1.5 mg/mL in Blocking Buffer) for 600s. Following loading, sensors were incubated for 420s in Blocking Buffer prior to binding assessment of the Fabs. Association of Fabs was measured for 300 s and dissociation was measured for up to 3,600 s in Blocking Buffer. All the assays were performed with agitation set to 1,000 rpm at 30 °C. Data analysis and curve fitting were carried out using Octet analysis software, version 11-12. Experimental data were fitted using a 1:1 binding model. Global analyses of the complete data sets assuming binding was reversible (full dissociation) were carried out using nonlinear least-squares fitting allowing a single set of binding parameters to be obtained simultaneously for all concentrations used in each experiment.

Cryo-EM specimen preparation and data collection.

A sample of SARS CoV-2 spike in PBS was mixed with Fab A23-58.1 at a molar ratio of 1.2 Fab per protomer. The final protein concentration was 0.5 mg/ml. n-Dodecyl β-D-maltoside (DDM) detergent was added shortly before vitrification to a concentration of 0.005%. Quantifoil R 2/2 gold grids were subjected to glow discharging in a PELCO easiGlow device (air pressure: 0.39 mBar, current: 20 mA, duration: 30 s) immediately before specimen preparation. Cryo-EM grids were prepared using an FEI Vitrobot Mark IV plunger with the following settings: chamber temperature of 4°C, chamber humidity of 95%, blotting force of -5, blotting time of 3 s, and drop volume of 2.7 µl. Datasets were collected at the National CryoEM Facility (NCEF), National Cancer Institute, on a Thermo Scientific Titan Krios G3 electron microscope equipped with a Gatan Quantum GIF energy filter (slit width: 20 eV) and a Gatan K3 direct electron detector (Table S2). Four movies per hole were recorded in the counting mode using Latitude software. The dose rate was 14.65 e-/s/pixel.

Cryo-EM data processing and model fitting

Data process workflow, including Motion correction, CTF estimation, particle picking and extraction, 2D classification, ab initio reconstruction, homogeneous refinement, heterogeneous refinement, non-uniform refinement, local refinement and local resolution estimation, were carried out with C1 symmetry in cryoSPARC 2.15 (47) For local refinement to resolve the RBD-antibody interface, a mask for the entire spike-antibody complex without the RBD-antibody region was used to extract the particles and a mask encompassing the RBD-antibody region was used for refinement. The overall resolution was 3.39 Å for the map of antibody-bound spike, 3.89 Å for the map of RBD:antibody after local refinement. The coordinates for the SARS-CoV-2 spike with three ACE2 molecules bound at pH 7.4 (PDB ID: 7KMS) were used as initial models for fitting the cryo-EM map. Iterative manual model building and real space refinement were carried out in Coot (48) and in Phenix (49), respectively. Molprobity (50) was used to validate geometry and check structure quality at each iteration step. UCSF Chimera and ChimeraX were used for map fitting and manipulation (51).

Negative-stain electron microscopy.

Protein samples were diluted to a concentration of approximately 0.02 mg/ml with 10 mM HEPES, pH 7.4, supplemented with 150 mM NaCl. A 4.8-µl drop of the diluted sample was placed on a freshly glow-discharged carbon-coated copper grid for 15 s. The drop was then removed with filter paper, and the grid was washed with three drops of the same buffer. Protein molecules adsorbed to the carbon were negatively stained by applying consecutively three drops of 0.75% uranyl formate, and the grid was allowed to air-dry. Datasets were collected using a Thermo Scientific Talos F200C transmission electron microscope operated at 200 kV and equipped with a Ceta camera. The nominal magnification was 57,000x, corresponding to a pixel size of 2.53 Å, and the defocus was set at -1.2 µm. Data was collected automatically using EPU. Single particle analysis was performed using CryoSPARC (47).

Selection of rcVSV SARS CoV-2 virus escape variants using monoclonal antibodies

A replication competent vesicular stomatitis virus (rcVSV) with its native glycoprotein replaced by the Wuhan-1 spike protein (rcVSV SARS CoV-2) that contains a 21 amino acid deletion at the C-terminal region (33) (generous gift of Kartik Chandran and Rohit Jangra). Passage 7 virus was passaged twice on Vero cells to obtain a polyclonal stock. A single plaque from this 9$^{th}$ passage was double plaque purified and expanded on Vero cells to create monoclonal virus population. The reference genome for this stock was sequence using Illumina-based sequencing as described below.

To select for virus escape variants, an equal volume of clonal population of rcVSV SARS CoV-2 was mixed with serial dilutions of antibodies (5-fold) in DMEM supplemented with 10% FCS and Glutamine to give an MOI of 0.1 - 0.001 at the desired final antibody concentration (range 5.1e-6 to 50 mg/ml and 0 mg/ml). Virus:antibody mixtures were incubated at room temperature for 1 hour. After incubation, 300 µl of virus:antibody mixtures were added to 1 x $10^5$ Vero E6 cells in 12 well plates for 1 hour at $37^0$C, 5% $CO_2$. The plates were rotated every 15 minutes to prevent drying. After absorption, 700 µl of additional antibodies mixture was added to each well at their respective concentration. Cells were incubated for 72hrs at $37^0$C, 5% $CO_2$. Virus replication was monitored using cytopathic effect and supernatant was collected from the wells with cytopathic effect. Harvested supernatant was clarified by centrifugation at 3750rpm for 10 minutes. For the subsequent rounds of selection, clarified supernatant from the well with the highest concentration of antibody that has CPE >20% supernatant was diluted prior to being mixed with equal volume of antibodies as in the initial round of selection. Infection, monitoring and collection of supernatants was performed as in the initial round.

Shotgun sequencing of rcVSV SARS CoV2 supernatants

Total RNA was extracted from clarified supernatants using QIAmp viral RNA mini extraction kit (Qiagen) following the manufacturer's recommended protocol. Purified RNA was fragmented using NEBNext Ultra II RNA Library Prep reagents, then reverse transcribed using random hexamers, and double-stranded cDNA was synthesized (New England BioLabs) as previously described (52). Double-stranded cDNA was purified using magnetic beads (MagBio Genomics) and barcoded Illumina-ready libraries were subsequently prepared (New England BioLabs). The libraries were sequenced as paired-end 2x150 base pair NextSeq 2000 reads.

Spike SNP variant calls of rcVSV antibody induced revertants

Raw sequencing reads were demultiplexed and trimmed to remove adaptor sequences and low quality bases. They were then aligned against the reference viral genome with Bowtie (v2.4.2). Single nucleotide polymorphisms (SNPs) were called using HaplotypeCaller from the Genome Analysis Tool Kit (GATK, v4.1.9.0). The HaplotypeCaller parameter, "--sample-ploidy", was set to 100 in order to identify SNPs with a prevalence of at least 1%. SNPs for all samples were then aggregated, interrogated and translated using custom scripts. A SNP and correlated amino acid translation for the spike protein was considered positive if it was present at a frequency of greater than 0.1 (10%) and showed an increasing frequency from round 1 to round 2 of the antibody selections.

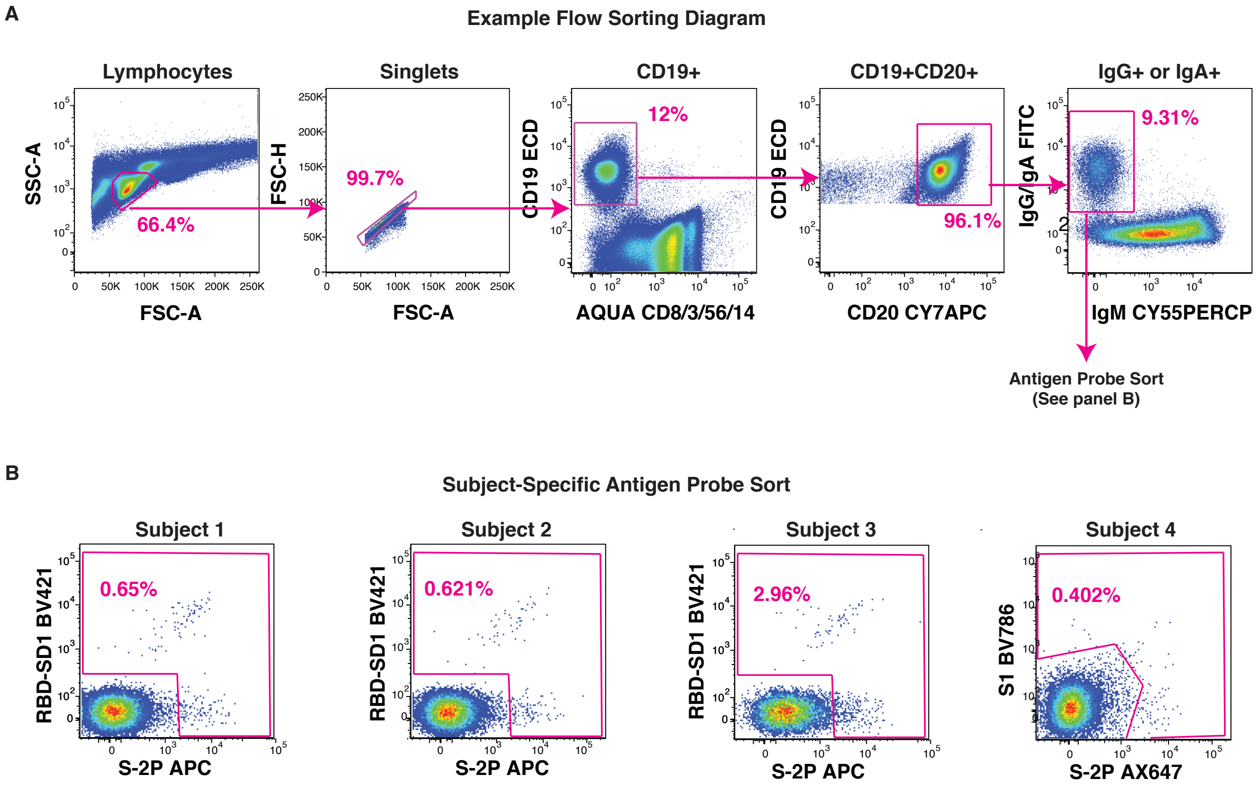

**Fig. S1. Example flow sorting schematic and subject-specific antigen sort gates.**

**(A)** Flow cytometry sorting scheme for subject PBMCs in order to isolate antigen positive CD19+/CD20+//IgM-/IgG+ or IgA+ B cells.

**(B)** B cells that are single or dual positive for sorted using the indicated gates and probes.

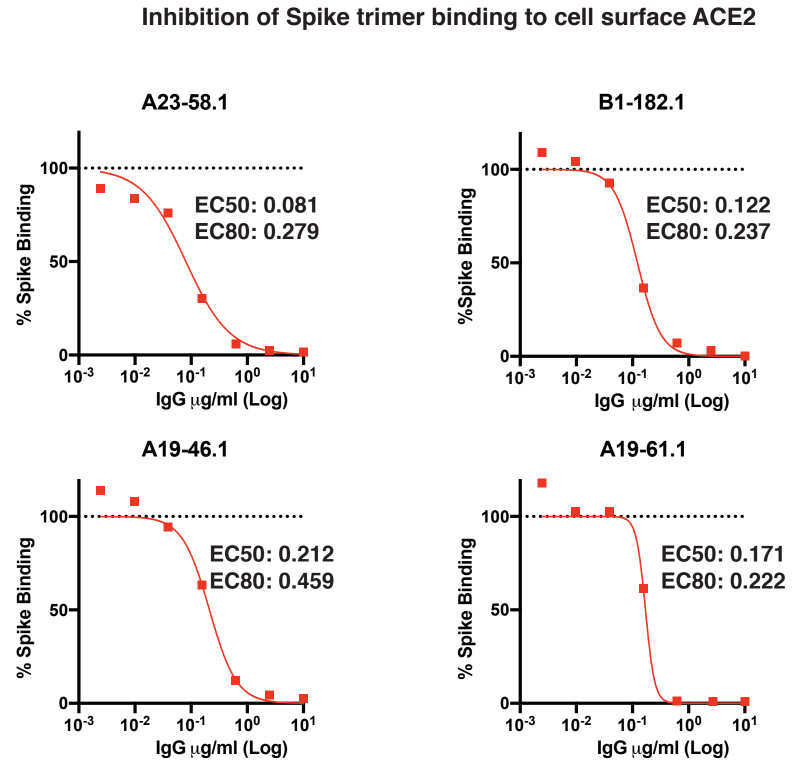

**Fig. S2. Inhibition of Spike trimer binding to cell surface ACE2.**

The indicated antibodies were mixed with Spike trimer at increasing concentrations of antibody prior to incubating with cells expressing ACE2. Spike binding was determined using flow cytometric analysis. Shown are the relative binding percentage as a function of antibody concentration.

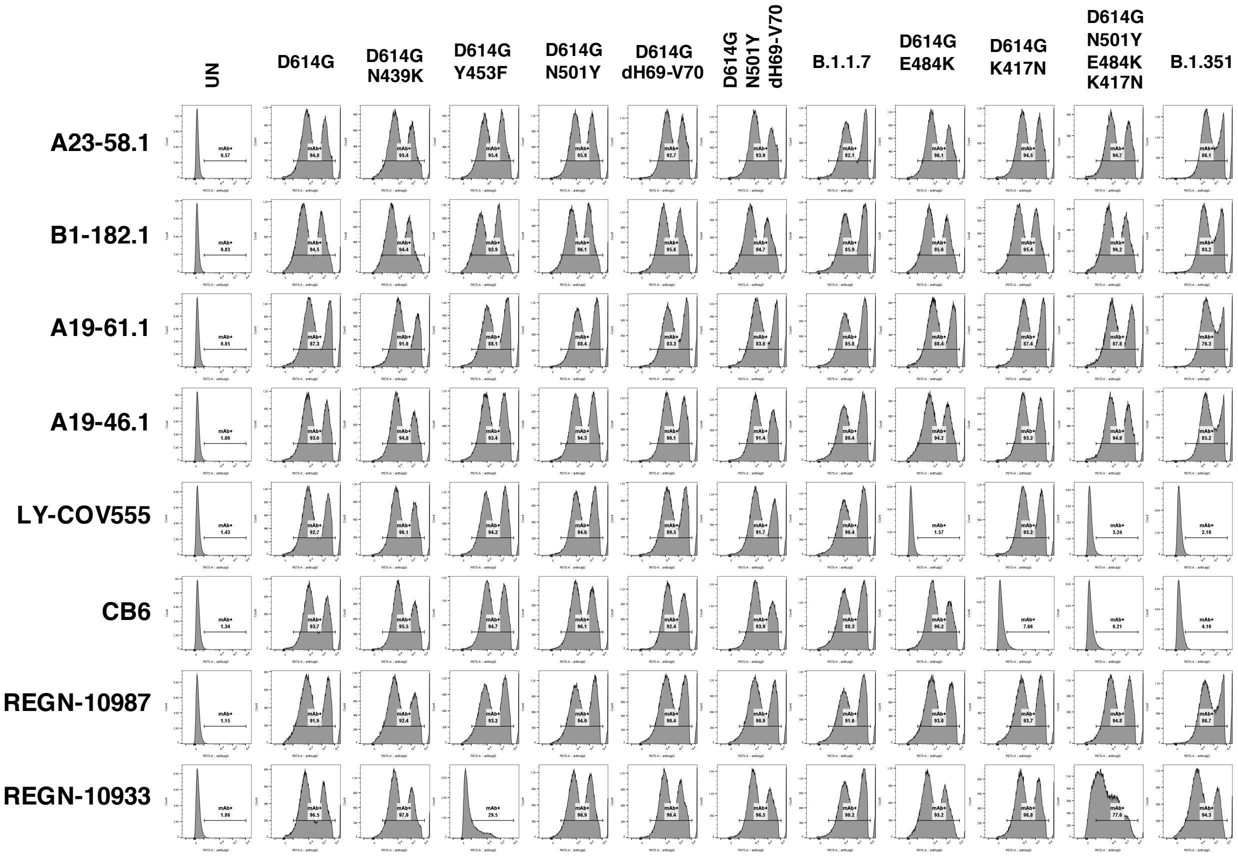

**Fig. S3. Binding of antibodies to cell surface expressed natural variant Spikes.**

The indicated antibodies were incubated with cells expressing the indicated natural spike variants and binding was determined using flow cytometric analysis. Shown are histogram plots for each antibody-spike protein combination.

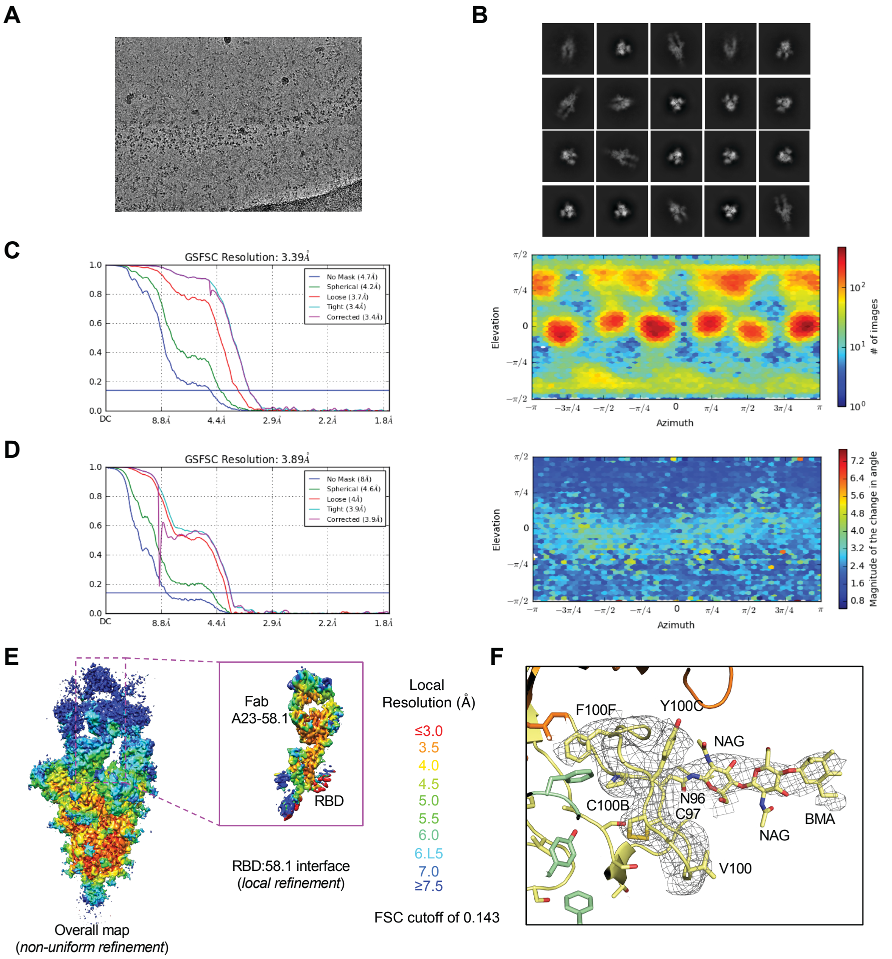

**Figure S4. Cryo-EM details of A23-58.1 Fab in complex with SARS-CoV-2 HexaPro spike.**
**(A)** Representative micrograph.
**(B)** Representative 2D class averages.
**(C)** The gold-standard Fourier shell correlation resulted in a resolution of 3.39 Å for the overall map using non-uniform refinement with C1 symmetry (left panel); the orientations of all particles used in the final refinement are shown as a heatmap (right panel).
**(D)** The gold-standard Fourier shell correlation resulted in a resolution of 3.89 Å for the masked local refinement of the RBD:A23-58.1 interface (left panel) obtained using particle subtraction followed by local refinement; the orientations of all particles used in the local refinement are shown as a heatmap (right panel).
**(E)** The local resolution of the final overall map and locally refined map is shown contoured at 0.068 (4.5s) and 0.105 (13.6s), respectively. Resolution estimation was generated through cryoSPARC using an FSC cutoff of 0.143.
**(F)** Representative density is shown for the CDR H3 loop of A23-58.1. The disulfide bond between Cys97 and Cys100B and glycan on Asn96 are defined. Antibody residue carbon atoms are colored in yellow, oxygen in red, nitrogen in blue; RBD is colored in palegreen and light chain is in orange. The contour level is 1.5s.

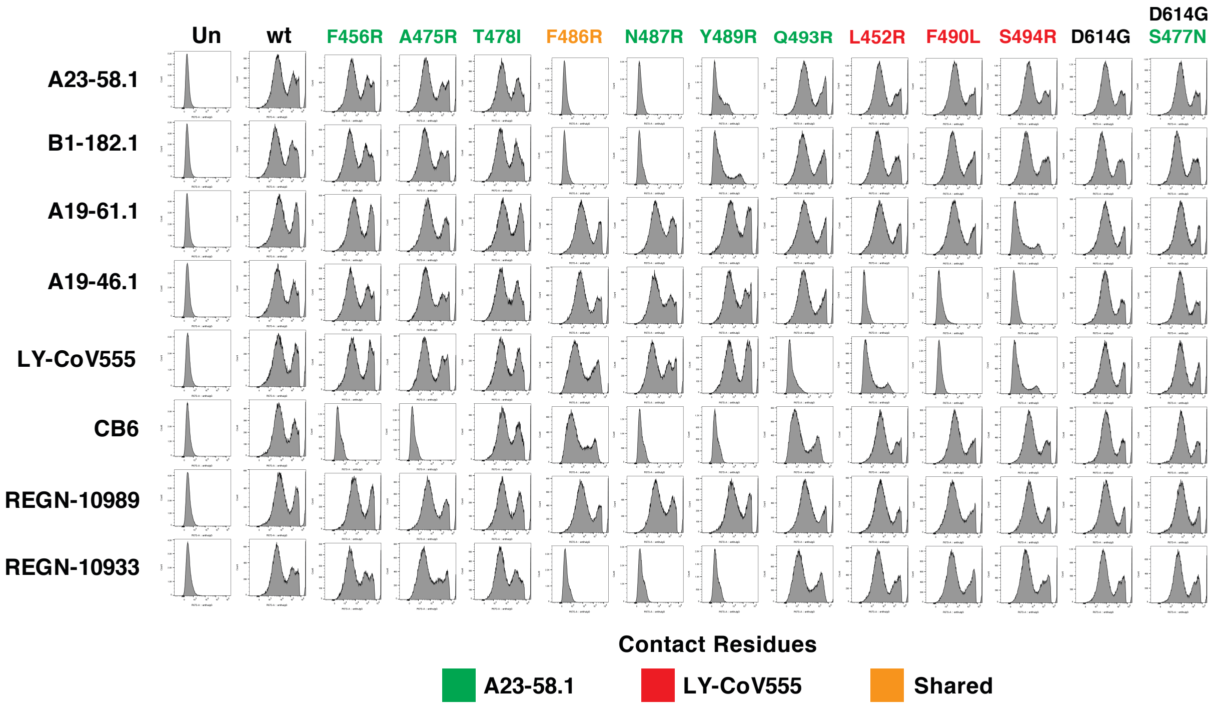

**Fig. S5. Binding of antibodies to cell surface expressed Spike mutants.**

The indicated antibodies were incubated with cells expressing the indicated spike mutation(s) and binding was determined using flow cytometric analysis. Shown are histogram plots for each antibody-spike protein combination.

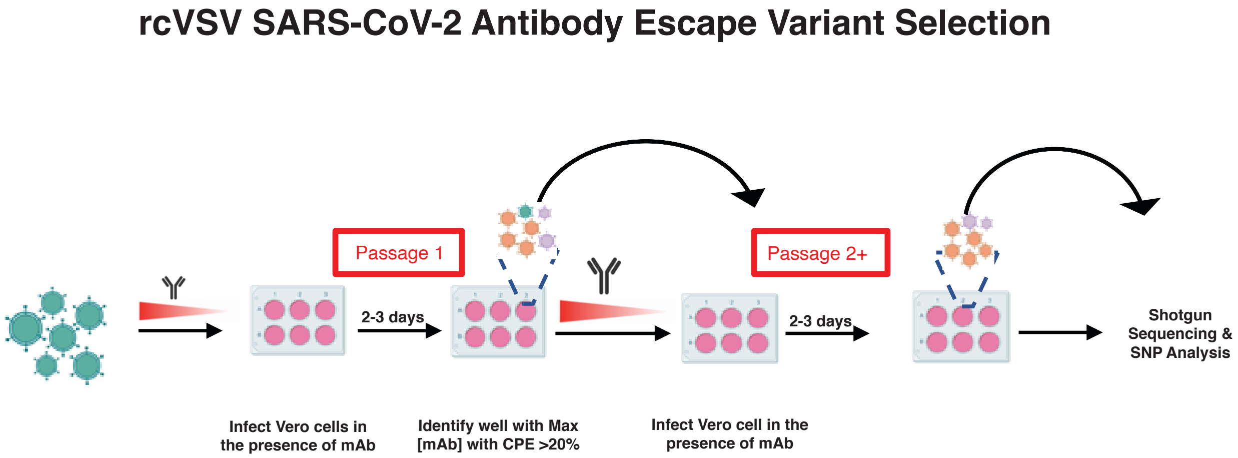

**Fig. S6. Schematic of rcVSV selection of escape variants to monoclonal antibodies.**

Virus and antibody mixtures are incubated at decreasing concentrations on Vero cells and cytopathic effect (CPE) determined in each well after 48-72 hours. The well at the highest concentration with at least 20% CPE was passaged forward into a second round where it is incubated in the presence of the same antibody concentration s as in round 1 and assessed for CPE as previously. Subsequent rounds can be carried forward as desired or harvested supernatants can be evaluated for escape variant sequences using shotgun sequencing and SNP variant call analyses.

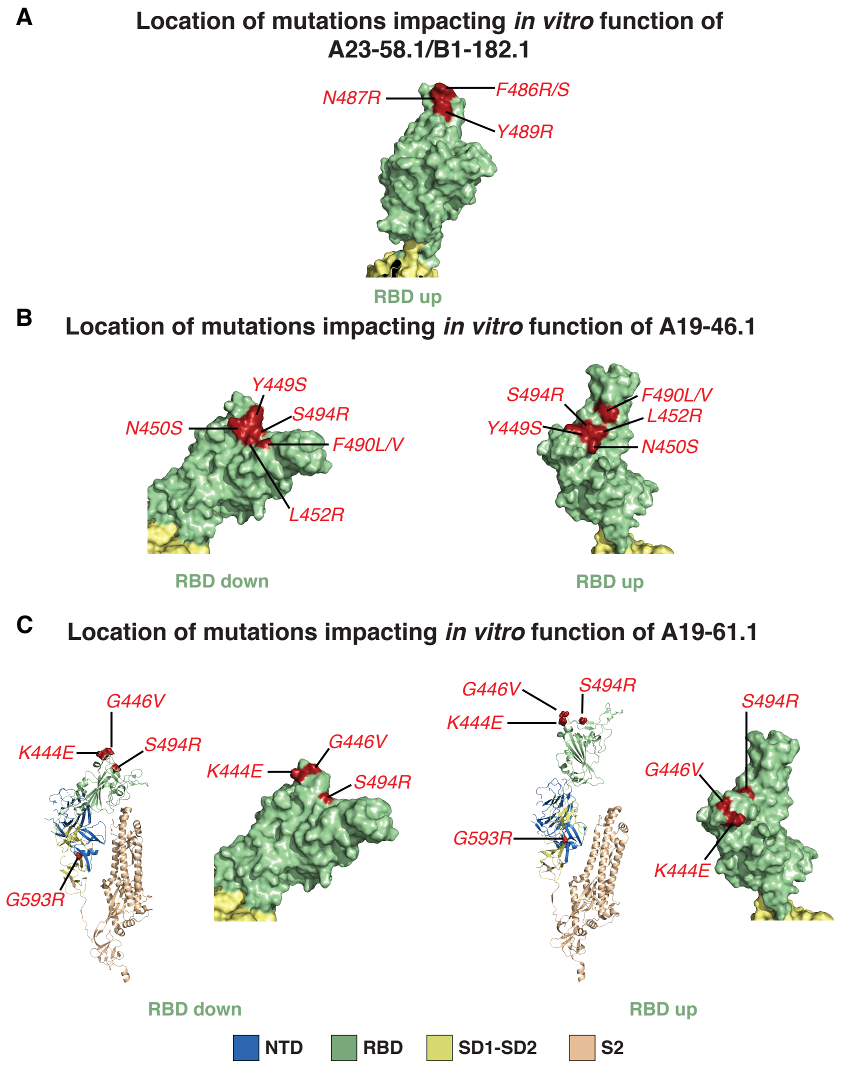

**Figure S7. Location of mutations impacting function of A23-58.1, B1-182.1, A19-46.1 and A19-61.1**

**(A)** Location of mutations identified impacting binding or neutralizing activity of A23-58.1 and B1-182.1 shown on model of RBD in the up position.

**(B)** Location of mutations identified impacting binding or neutralizing activity of A19-46.1 shown on model of RBD in the down (left) or up (right) position.

**(C)** Location of mutations identified impacting binding or neutralizing activity on A19-61.1 shown on single spike protomer or RBD models. Models are shown with RBD in the down (left) or up (right) position.

All models in rendered in Pymol using PDB:6Z97.

### Gene Family Usage

| Name | VH | DH | JH | CDRH3 | VL | JL | CDRL3 |
|---|---|---|---|---|---|---|---|
| **A19-46.1** | IGHV3-30*03, IGHV3-30-5*01 | IGHD1-36*01 | IGHJ6*02 | CARGWAYWELLPDYYYGMDVW | IGLV8-61*01 | IGLJ3*01, IGLJ2*01 | CVLYMGRGIVVF |
| **A19-61.1** | IGHV3-30-3*01, IGHV3-30*17 | IGHD6-19*01 | IGHJ6*02 | CARDLAIAVAGTWHYYNGMDVW | IGKV1D-12*02, IGKV1-12*02 | IGKJ5*01 | CQQAKSFPITF |
| **A23-58.1** | IGHV1-58*01 | IGHD2-8*01 | IGHJ6*02 | CAAPNCSNVVCYDGFDIW | IGKV3-20*01 | IGKJ1*01 | CQQYGTSPWTF |
| **B1-182.1** | IGHV1-58*01 | IGHD2-15*01 | IGHJ3*02 | CAAPYCSGGSCFDGFDIW | IGKV3-20*01 | IGKJ1*01 | CQQYGNSPWTF |

**Table S1.** Genetic origins were assigned using SONAR. In cases where multiple closely related genes produced indistinguishable alignments with the query sequence, all possibilities are shown.

**Table S2. Cryo-EM Data Collection, Refinement and Validation Statistics for A23-58.1 in Complex with Spike**

| | SARS-CoV-2 spike in complex with A23-58.1 (EMD-23499) (PDB 7LRT) | | A23-58.1:RBD complex after local refinement (EMD-23498) (PDB 7LRS) |
|---|---|---|---|
| **Data collection and processing** | | | |
| Magnification | | 105,000 | |
| Voltage (kV) | | 300 | |
| Electron exposure ($e^{-}$/Å$^2$) | | 40.0 | |
| Defocus range (μm) | | -1.1 to -2.1 | |
| Pixel size (Å) | | 0.855 | |
| Symmetry imposed | | C1 | |
| Final particle images (no.) | | 123,016 | |
| Map resolution (Å) | 3.47 | | 3.89 |
| FSC threshold | 0.143 | | 0.143 |
| **Refinement** | | | |
| Initial model used (PDB code) | | 7KMS | |
| Model resolution (Å) | | 3.64 | |
| FSC threshold | 0.143 | | 0.143 |
| Map sharpening *B* factor (Å$^2$) | -71.1 | | -45.7 |
| Model composition | | | |
| Non-hydrogen atoms | 33465 | | 4912 |
| Protein residues | 4152 | | 625 |
| Ligands | 84 | | 8 |
| *B* factors (Å$^2$)(mean) | | | |
| Protein | 186 | | 78 |
| Ligand | 189 | | 104 |
| R.m.s. deviations | | | |
| Bond lengths (Å) | 0.004 | | 0.003 |
| Bond angles (°) | 0.623 | | 0.644 |
| Validation | | | |
| MolProbity score | 1.94 | | 2.05 |
| Clash score | 9.07 | | 8.70 |
| Poor rotamers (%) | 0.0 | | 0.19 |
| Ramachandran plot | | | |
| Favored (%) | 92.8 | | 88.7 |
| Allowed | 6.9 | | 10.7 |
| Disallowed | 0.3 | | 0.6 |

**Table S3. Interactions between A23-58.1 and the spike**

| Binding area of the epitope and paratope | | | |
|---|---|---|---|
| | Paratope (Å$^2$) | | Epitope (Å$^2$) |
| Heavy chain | 437.7 | | 455.6 |
| Light chain | 163.6 | | 151.1 |
| Hydrogen bonds | | | |
| | Antibody | Distance (Å) | RBD |
| 1 | D:CYS 100B[ N ] | 3.17 | C:ALA 475[ O ] |
| 2 | D:ASP 100D[ OD1] | 3.71 | C:SER 477[ N ] |
| 3 | D:ASP 100D [ OD2] | 3.73 | C:SER 477[ N ] |
| 4 | D:ASP 100D [ OD1] | 2.53 | C:ASN 487[ ND2] |
| 5 | E:TYR 32[ OH ] | 3.81 | C:THR 478[ OG1] |

41. J. S. McLellan, M. Pancera, C. Carrico, J. Gorman, J.-P. Julien, R. Khayat, R. Louder, R. Pejchal, M. Sastry, K. Dai, S. O'Dell, N. Patel, S. Shahzad-ul-Hussan, Y. Yang, B. Zhang, T. Zhou, J. Zhu, J. C. Boyington, G.-Y. Chuang, D. Diwanji, I. Georgiev, Y. Do Kwon, D. Lee, M. K. Louder, S. Moquin, S. D. Schmidt, Z.-Y. Yang, M. Bonsignori, J. a Crump, S. H. Kapiga, N.

E. Sam, B. F. Haynes, D. R. Burton, W. C. Koff, L. M. Walker, S. Phogat, R. Wyatt, J. Orwenyo, L.-X. Wang, J. Arthos, C. a Bewley, J. R. Mascola, G. J. Nabel, W. R. Schief, A. B. Ward, I. a Wilson, P. D. Kwong, Structure of HIV-1 gp120 V1/V2 domain with broadly neutralizing antibody PG9. Nature. 480, 336–43 (2011)

42. D. H. Barouch, Z. Yang, W. Kong, B. Korioth-Schmitz, S. M. Sumida, D. M. Truitt, M. G. Kishko, J. C. Arthur, A. Miura, J. R. Mascola, N. L. Letvin, G. J. Nabel, A Human T-Cell Leukemia Virus Type 1 Regulatory Element Enhances the Immunogenicity of Human Immunodeficiency Virus Type 1 DNA Vaccines in Mice and Nonhuman Primates. J. Virol. 79, 8828–8834 (2005).

43. A. T. Catanzaro, M. Roederer, R. A. Koup, R. T. Bailer, M. E. Enama, M. C. Nason, J. E. Martin, S. Rucker, C. A. Andrews, P. L. Gomez, J. R. Mascola, G. J. Nabel, B. S. Graham, VRC 007 Study Team, Phase I clinical evaluation of a six-plasmid multiclade HIV-1 DNA candidate vaccine. Vaccine. 25, 4085–92 (2007).

44. L. Naldini, U. Blomer, F. H. Gage, D. Trono, I. M. Verma, Efficient transfer, integration, and sustained long-term expression of the transgene in adult rat brains injected with a lentiviral vector. Proc. Natl. Acad. Sci. U. S. A. 93, 11382–11388 (1996).

45. Z. Y. Yang, H. C. Werner, W. P. Kong, K. Leung, E. Traggiai, A. Lanzavecchia, G. J. Nabel, Evasion of antibody neutralization in emerging severe acute respiratory syndrome coronaviruses. Proc. Natl. Acad. Sci. U. S. A. 102, 797–801 (2005).

46. L. Wang, W. Shi, M. G. Joyce, K. Modjarrad, Y. Zhang, K. Leung, C. R. Lees, T. Zhou, H. M. Yassine, M. Kanekiyo, Z. Y. Yang, X. Chen, M. M. Becker, M. Freeman, L. Vogel, J. C. Johnson, G. Olinger, J. P. Todd, U. Bagci, J. Solomon, D. J. Mollura, L. Hensley, P. Jahrling, M. R. Denison, S. S. Rao, K. Subbarao, P. D. Kwong, J. R. Mascola, W. P. Kong, B. S. Graham,

Evaluation of candidate vaccine approaches for MERS-CoV. Nat. Commun. 6 (2015), doi:10.1038/ncomms8712.
